# Zygotic genome activation by the totipotency pioneer factor Nr5a2

**DOI:** 10.1101/2022.05.17.492379

**Authors:** Johanna Gassler, Wataru Kobayashi, Imre Gáspár, Siwat Ruangroengkulrith, Maximilian Kümmecke, Pavel Kravchenko, Maciej Zaczek, Antoine Vallot, Laura Gomez Hernandez, Laura Cuenca Rico, Sabrina Ladstätter, Kikuë Tachibana

## Abstract

Life begins with a switch in genetic control from the maternal to the embryonic genome during zygotic genome activation (ZGA) in totipotent embryos. Despite its importance, the essential regulators of ZGA remain largely unknown in mammals. Based on *de novo* motif searches, we identified the orphan nuclear receptor Nr5a2 as a key activator of major ZGA in mouse embryos. Nr5a2 binds to its motif within a subtype of *SINE B1/Alu* transposable elements found in *cis*-regulatory regions of ZGA genes. Chemical inhibition suggests that 72% of ZGA genes are regulated by Nr5a2 and potentially other orphan nuclear family receptors. Consistent with a role in ZGA, Nr5a2 is required for progression beyond the 2-cell stage. Nr5a2 promotes chromatin accessibility during ZGA and binds to entry/exit sites of nucleosomal DNA *in vitro*. We conclude that Nr5a2 is an essential pioneer factor that distinctly regulates totipotency and pluripotency during mammalian development.

**One-Sentence Summary:** Nr5a2 is an essential pioneer transcription factor that activates expression of zygotic genes in mouse embryos.

## Main Text

Since the oocyte cytoplasm is sufficient to reprogram somatic cell nuclei to totipotency, it is thought that the factors responsible for reprogramming and triggering ZGA in the early embryo are maternally provided (*1*). The transcriptional “awakening” of the embryonic genome occurs in at least two waves, minor ZGA in the zygote and major ZGA (hereafter referred to as ZGA) in the 2-cell embryo of mice (Fig. S1A) (*2, 3*). Embryonic transcription is required for progression beyond the 2-cell stage (*4*).

The transcription factors that initiate ZGA appear to be poorly conserved between species. During *Drosophila* embryogenesis, transcription factors including the diptera-specific Zelda, GAGA factor (GAF) and Chromatin-linked adaptor for male-specific lethal protein (CLAMP) are essential for ZGA (*5-7*). In zebrafish and frogs, pluripotency factors belonging to the POU and Sox families and Nanog are required for ZGA (*8-10*). In contrast, the pluripotency factor Oct4 is not essential for murine ZGA (*11*). Nfya, Yap1, Dux, Rarg, Dppa2 and Dppa4 have been linked to mammalian ZGA (*12-17*) but are not required for progression beyond the 2-cell embryo in knockdown or genetic knockout models (*12, 18-22*). The essential transcription factors that activate mammalian embryonic genomes therefore remain unknown.

Transcription factors bind to regulatory elements in the genome to control gene expression and cell fate. Genomic DNA is wrapped around a histone octamer in the nucleosome, which is a barrier to transcription factor occupancy. Pioneer transcription factors (hereafter referred to as pioneer factors) are a class of transcription factors with the capacity to bind their (partial) motif on nucleosomal DNA *in vitro* and to be recruited to closed chromatin *in vivo*, eliciting local chromatin opening by diverse mechanisms (*23*). Pioneer factors cooperatively open the fruitfly and zebrafish genomes during ZGA (*6-9, 24, 25*). Whether pioneer activities initiate mammalian ZGA is not known. Here, we identified the orphan nuclear receptor Nr5a2 as a *bona fide* pioneer factor that activates genome-wide zygotic gene expression in mouse embryos.

### Nr5a2 is required for early embryonic development

We hypothesized that motifs of transcription factors regulating ZGA are enriched in the *cis*-regulatory regions of ZGA genes. To define ZGA genes, we compared transcription profiles of two mouse strains during the oocyte-to-embryo transition (OET). A total of 2508 “extended” ZGA genes were upregulated >4-fold in 2-cell embryos of the two strains. We classified the 985 genes that were common to both as “core” ZGA genes (Fig. S1B-E). Using a *de novo* motif search, we found a consensus sequence, comprising six motifs, that is enriched upstream of ∼70% of core and 77% of extended ZGA genes (compared to 46% non-ZGA genes) (Fig.1A and S1F). The sequence has 90% similarity to the *Short Interspersed Nuclear Element* (*SINE*) *B1* family of retrotransposons, which are related to the human *Alu* family, and contains variable pyrimidine/purine (YR) stretches (Fig 1A). A particular variant, CA, of the fifth YR stretch (YR5) correlates with higher chromatin accessibility and histone acetylation during major ZGA, implying a functional relevance. YR5 is located in motif 1, which shows a high frequency and mean occurrence upstream of ZGA genes (Fig. S1F). The CA version of motif 1 contains the consensus sequences for orphan nuclear receptors Nr5a2 (T**CA**AGG<u>C</u>CA, hereafter Nr5a2 motif) and Esrrb (T**CA**AGG<u>T</u>CA, hereafter Esrrb motif) and nuclear hormone receptor “retinoic acid receptor gamma” (Rarg) (AGGT**CA**AGGTCA) (*26, 27*). Given that Rarg is not essential for ZGA since *Rarg*^*-/-*^ females are fertile (*18*), we focused on the orphan nuclear receptors. *Esrrb*^*-/-*^ and *Nr5a2*^*-/-*^ embryos produced from heterozygote intercrosses die around implantation (*28, 29*). It is therefore unknown whether these transcription factors have functions during ZGA.

**Figure 1.**
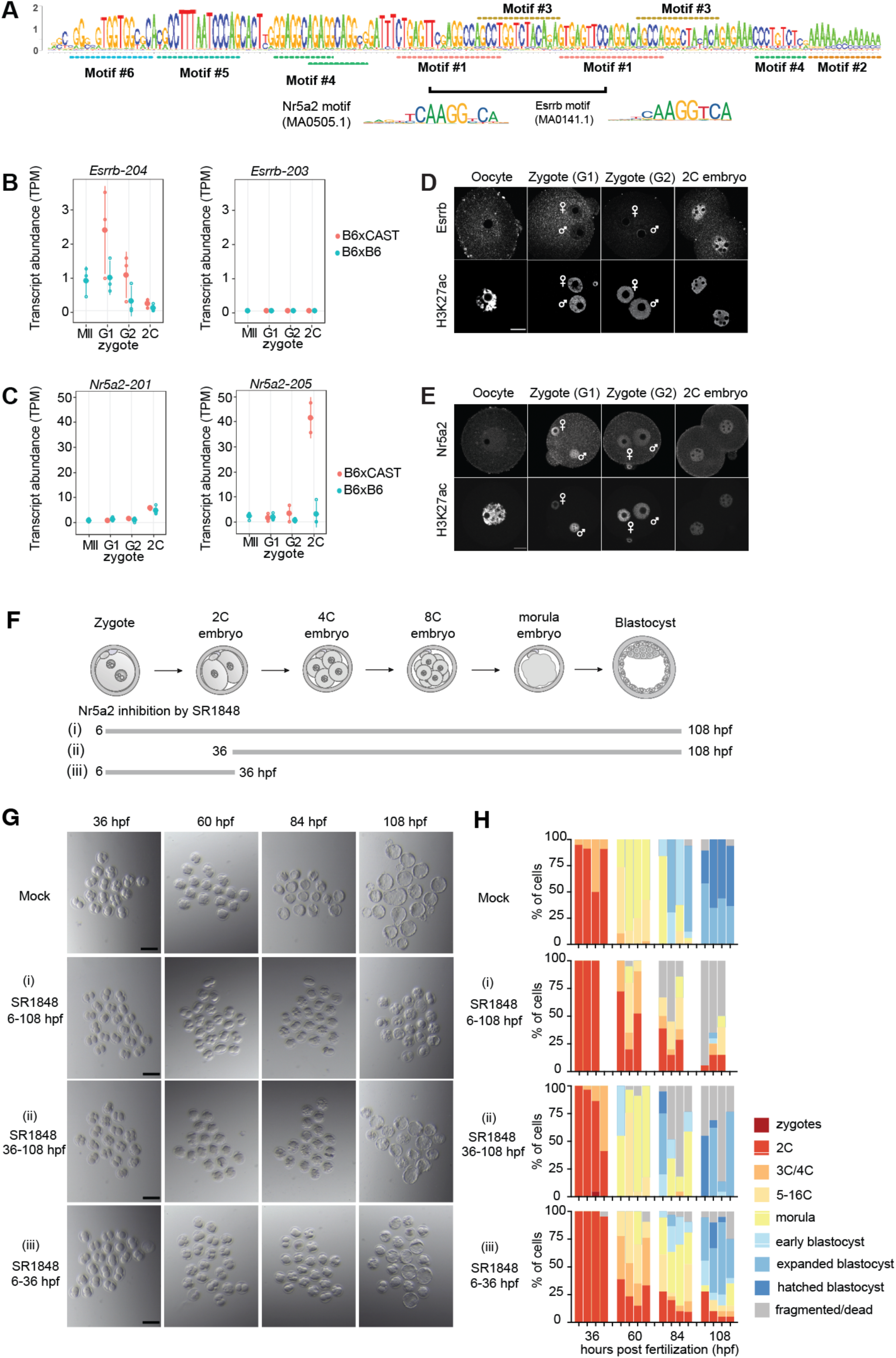
Nr5a2 is required for early embryonic development. (**A**) Sequence logo of the identified supermotif with the consisting motifs highlighted. Motif #1 resembles the cognate binding sequence of Nr5a2 and Esrrb. (**B and C**) Transcript abundance of main protein coding isoforms of Esrrb (Esrrb-203 and Esrrb-204) (B) and Nr5a2 (Nr5a2-201 and Nr5a2-205) (C) during the oocyte to 2-cell (2C) embryo transition from pure B6 (B6xB6) and B6CASTF1 (B6xCAST) mice (see methods). (**D and E**) Representative immunofluorescence images depicting Esrrb (D) and Nr5a2 (E) in embryonic stages. Maternal and paternal zygotic nuclei are indicated by the symbols. H3K27ac immunofluorescence is shown to indicate proper antibody penetration and to outline the nuclei. Depicted are single z-slices. Scale bar represents 20 μm. (**F**) Schematic of early embryo stages including timing and length of inhibitor treatment. (**G**) Stereomicroscopic example images of the embryonic stages observed to the indicated time points in the different conditions. Mock control embryos were treated with the same volume of DMSO as inhibitors. Scale bars are 150 μm. (**H**) Quantification of four replicate experiments of embryonic development. Sample sizes are: mock: n= 19, 23, 16, 33; SR1848 (6-108 hpf): n= 18, 20, 21; SR1848 (6-36 hpf): n=18, 20, 20, 30 cells; SR1848 (36-108 hpf): n=20, 21, 17, 29; each experiment comprising of 6-12 females.

To assess whether Nr5a2 and Esrrb are ZGA regulators, we first examined their expression during the OET. Nr5a2 and Esrrb transcript isoforms were present in oocytes and upregulated in 2-cell embryos (Fig. 1B and 1C). Esrrb protein was detected by immunofluorescent staining in 2-cell embryos but not earlier (Fig. 1D). In contrast, Nr5a2 protein was detected in oocytes, zygotes and 2-cell embryos (Fig. 1E). Since Nr5a2 protein is present in embryos prior to ZGA, we focused on this transcription factor as a candidate ZGA regulator.

To determine if Nr5a2 has an early embryonic function that would be consistent with regulating ZGA, we isolated zygotes at 5 hours post fertilization (hpf) and examined their developmental potential over 4 days in culture (Fig. 1F-H, mock). We selectively inhibited Nr5a2 by treating embryos with the chemical compound SR1848, which does not inhibit the closely related Nr5a1 (*30, 31*). In contrast with control embryos, embryos treated from 6 hpf with SR1848 did not form blastocysts and were fragmented or dead at 108 hpf (Fig. 1G and 1H, scheme (i)). These findings are consistent with a function of Nr5a2 in maintaining naïve pluripotency in mouse ES cells (*26*). Intriguingly, embryos treated with SR1848 from 36 hpf (2-cell stage) had a less severe phenotype than those treated from 6 hpf (Fig. 1G and 1H, scheme (ii)), suggesting that Nr5a2 activity is required both before and after the 2-cell stage. Indeed, transient inhibition of Nr5a2 from 6-36 hpf caused either an immediate arrest at the 2-cell stage or a >24 h developmental delay (Fig. 1G and 1H, scheme (iii)). Overall, these findings suggest that Nr5a2 plays multiple roles in early development, including a hitherto unknown function between fertilization and the 2-cell stage, when ZGA occurs. Even though SR1848 is known to inhibit Nr5a2 and not Nr5a1, it is conceivable that it might have other targets. We therefore consider it possible that Nr5a2 functions redundantly with other members of the nuclear receptor family in regulating early development.

### Nr5a2 and Esrrb contribute to ZGA

To monitor ZGA directly, we examined nascent ZGA transcripts by single-molecule fluorescence *in situ* hybridization (ZGA-FISH). We designed for robustness two sets of FISH probes to detect ∼80 nascent ZGA transcripts each per fluorescence channel and used single-molecule FISH analysis algorithms to quantify *de novo* expression of ZGA genes (Fig. S1D) (*32, 33, 34*). We determined the lower threshold of the ZGA-FISH assay by culturing isolated zygotes (6 hpf) with triptolide to degrade RNA polymerase II. The total copy number of nuclear transcripts was reduced by ∼62% in triptolide-treated 34 hpf 2-cell embryos compared to DMSO-treated controls (Fig. S2A and S2B).

Culturing zygotes with SR1848 resulted in a dose-dependent reduction of nascent ZGA transcripts (∼20% and ∼40% for 5 μM and 10 μM SR1848, respectively) (Fig. 2A and 2B), suggesting that Nr5a2 and potentially other orphan nuclear receptors regulate ZGA. To test directly whether Nr5a2 is involved in ZGA, we microinjected oocytes with Nr5a2 siRNA, performed *in vitro* fertilization and analyzed 2-cell embryos by ZGA-FISH (Fig. 2C). Nr5a2 knockdown had a negligible effect on ZGA in 34 hpf 2-cell embryos (Fig. S2C and S2D). We considered that zygotic transcription of Nr5a2 might restore its function (Fig. 1C) and examined early 2-cell embryos (26 hpf) to minimize this potential effect. We found that Nr5a2 knockdown reduced nascent ZGA transcripts by ∼27% compared to controls (Fig. 2D). Expression of Nr5a2 mRNA under these conditions rescued ZGA, demonstrating specificity of the knockdown (Fig. 2E). Using a similar approach, we found that Esrrb knockdown repressed ZGA by ∼18% (Fig. 2D). These data suggest that Nr5a2 and Esrrb contribute to efficient ZGA.

**Figure 2.**
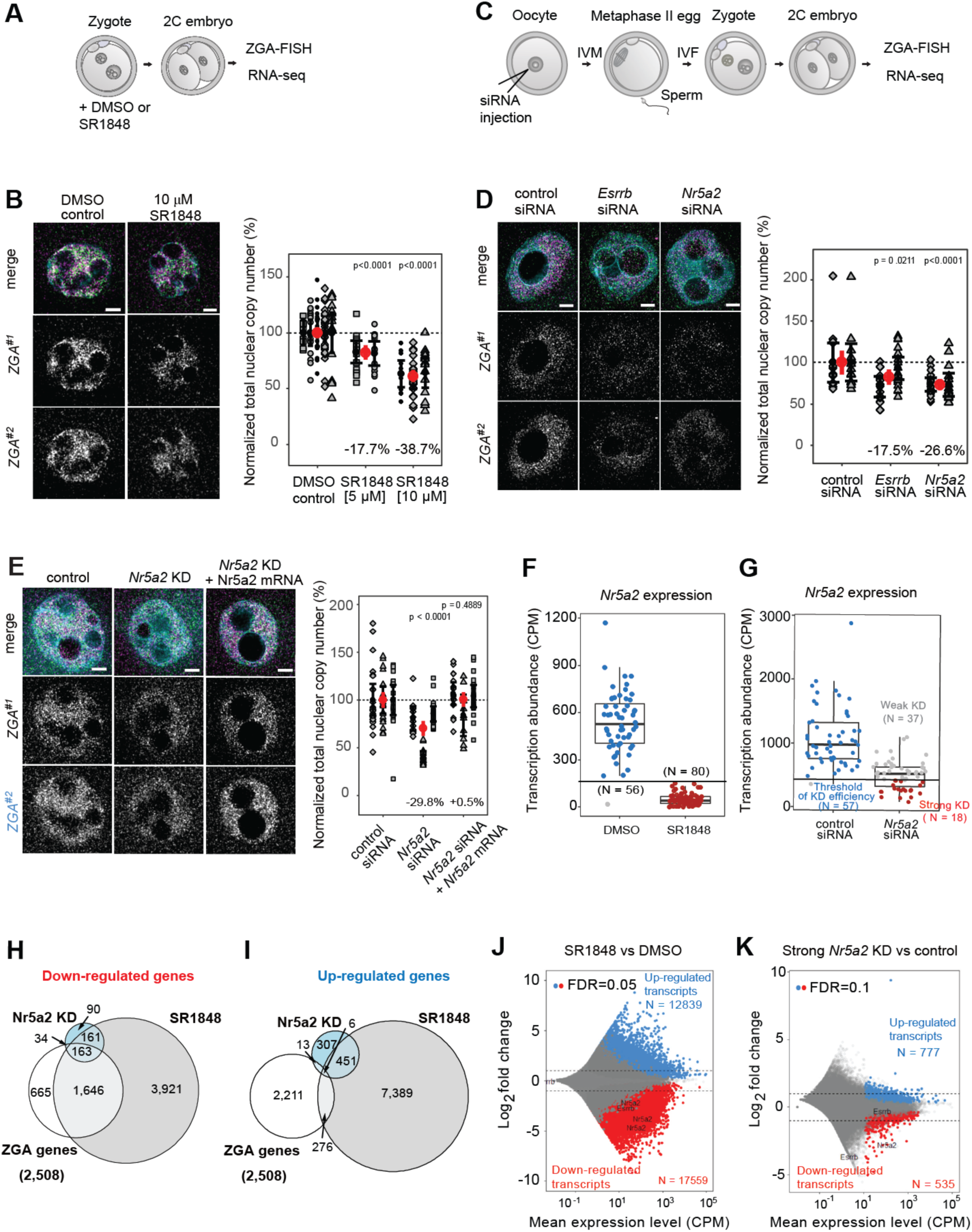
Nr5a2 and Esrrb are required for efficient ZGA. **(A)** Schematic illustration of experiments with DMSO control or SR1848 treated embryos. **(B)** Representative nascent ZGA-FISH images of SR1848 treated and DMSO control 2-cell embryos (34 hpf). Right panel shows a quantification of total nascent ZGA-FISH signal within nuclei of 2-cell embryos treated with different concentrations of SR1848 (5 or 10 μM) and their controls. Experimental replicates are shown with different bullet styles. Black dots and bars show the mean and 95% confidence interval per replicate, red dots and bars indicate the mean and 95% confidence interval of the experimental condition with all replicates merged (also for D and E). Number of nuclei analyzed in the replicates: control: n = 9, 18 and 19, 32, 18; 5 μM SR1848: n = 16, 17; 10 μM SR1848: n = 13, 25, 16. Scale bars are 5 μm. (**C**) Schematic illustration of experiments of siRNA knockdown in embryos. (**D**) Representative nascent ZGA-FISH images of *Esrrb, Nr5a2* knockdown embryos and their controls during early ZGA (26hpf). Right panel shows a quantification of the total nascent ZGA-FISH signal within nuclei of *Nr5a2, Esrrb* knockdown and control 2-cell embryos in early ZGA in two replicates. Number of nuclei analyzed in the replicates: control: n = 15, 15; *Esrrb* knockdown: n = 19, 16; *Nr5a2* knockdown: n = 16,18. Scale bars are 5 μm. (**E**) Representative nascent ZGA-FISH images of Nr5a2 knockdown, Nr5a2 knockdown supplemented with Nr5a2 mRNA and control embryos. Right panel shows a quantification of total nascent ZGA-FISH signal within nuclei of 2-cell embryos in the respective conditions. Number of nuclei analyzed in the replicates: control: n = 24, 20, 19; Nr5a2 knockdown: n = 13, 15, 20; Nr5a2 knockdown + Nr5a2 mRNA: n = 18, 11, 20. Scale bars are 5 μm. (**F**) Abundance of *Nr5a2* transcripts in 2-cell embryos treated by 10 μM SR1848 and their control as determined by single embryo RNA sequencing (Table S3). (**G**) Abundance of *Nr5a2* transcripts in *Nr5a2* knockdown 2-cell embryos and their control as determined by single embryo RNA sequencing (Table S4). (**H**) Euler diagrams showing the overlap of major ZGA genes, down-regulated genes in strong Nr5a2 knockdown and in SR1848-treated 2-cell embryos. (**I**) Euler diagrams showing the overlap of major ZGA genes, up-regulated genes in strong Nr5a2 knockdown and in SR1848-treated 2-cell embryos. (**J**) MA plot comparing chemically inhibited 2-cell embryos to control by DESeq2 (**K**) MA plot comparing strong Nr5a2 knockdown embryos to control by DESeq2.

To examine ZGA genome-wide, we performed single-embryo RNA-seq of 2-cell embryos at 34 hpf (Fig. 2A and 2C). Inhibition by SR1848 strongly decreased *Nr5a2* abundance (*31*) whilst the decrease was more moderate using siRNA knockdown at this time-point (Fig. 2F and 2G). We noticed that the knockdown efficiency varied between cells (Fig. 2G, S2E and S2F) and thus focused on embryos with the fewest Nr5a2 transcripts (Fig. 2G). Nr5a2 knockdown reduced the abundance of 197 out of 2,508 extended ZGA genes in 2-cell embryos, whereas culturing zygotes with SR1848 led to downregulation of 1,809, including most genes that were downregulated by Nr5a2 knockdown (Fig. 2H). By comparison, only 19 and 282 ZGA genes were upregulated by Nr5a2 knockdown and SR1848 treatment, respectively (Fig. 2I). Notably, SR1848 treatment reduced the transcript abundance of both Nr5a2 and Esrrb in 2-cell embryos, raising the possibility that the strong ZGA repression and compromised development are due to the combined loss of both transcription factors (Fig. 2J, 2K and S2G). We conclude that Nr5a2 is required for expression of major ZGA genes.

### Nr5a2 binds near TSS of Nr5a2-regulated ZGA genes

To determine whether Nr5a2 binds in the vicinity of ZGA genes in 2-cell embryos, we used a modified CUT&Tag approach (*35*) (Fig. 3A). Given that there are no precedents for mapping transcription factor binding sites in early mammalian embryos, we adapted CUT&Tag for ultra-low input samples, and verified the results with published data on histone modifications (Fig. S3A-C) (*36, 37*). We optimized conditions to examine Nr5a2 and Esrrb, which are both present in 2-cell embryos. We found that modified CUT&Tag of 2-cell embryos with Nr5a2 and Esrrb antibodies enriched for specific and overlapping genomic regions that were not observed with the IgG control (Fig. 3B, S3D and S3E). We identified 4,035 peaks enriched for both Nr5a2 and Esrrb, 4,524 peaks unique to Nr5a2 and 13,141 peaks unique to Esrrb (Fig. 3C).

**Figure 3.**
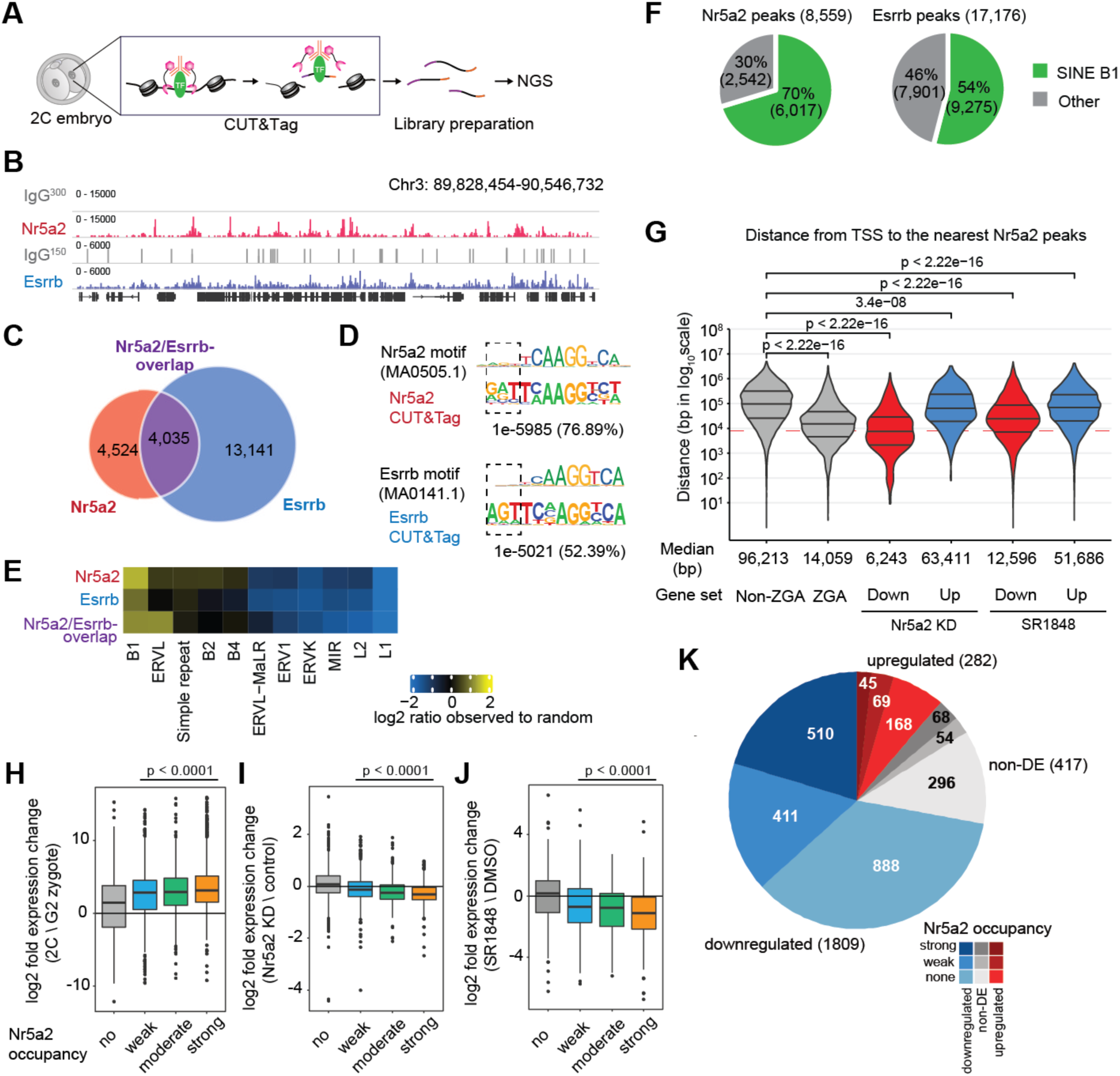
Genomic localization of Nr5a2 and Esrrb during ZGA. (**A**) Schematic illustration of CUT&Tag on 2-cell embryos. (**B**) Representative IGV snapshot shows enrichment of IgG control (grey), Nr5a2 (red) and Esrrb (blue) within the indicated region on chromosome 3. (**C**) Euler diagram showing overlap between Nr5a2 and Esrrb peaks in 2-cell embryos. (**D**) DNA sequence identified by Homer *de novo* motif analysis from Nr5a2 and Esrrb peaks in comparison to Nr5a2 (MA0505.1) and Esrrb (MA0141.1) motifs reported in the JASPAR database. The p-value of the motif comparison and percent of peaks containing the *de novo* motif are indicated. Black dot squares show three extended nucleotides which are identified from CUT&Tag in 2-cell embryos. (**E**) Heat maps show the enrichment of repeats (subfamily) in Nr5a2-unique, Esrrb-unique and Nr5a2/Esrrb-overlap peaks in 2-cell embryos. (**F**) Pie chart showing the percentage of peaks with *SINE B1*. (**G**) Violin plots showing the distance from TSS to the nearest Nr5a2 peaks between down-regulated and up-regulated genes in Nr5a2 KD embryos, ZGA genes and non-ZGA genes. Red dot line shows 8k bp. (**H-J**) Box plots showing the expression change of genes with no, weak, moderate or strong Nr5a2 CUT&Tag signal at Nr5a2 motifs in their 8k bp upstream regions (H) between G2 zygotes and 2-cell embryos, (I) in Nr5a2 and (J) KD in SR1848-treated 2-cell embryos. Bonferroni corrected p-values of pairwise Mann-Whitney U tests against genes with no Nr5a2 occupancy are shown. (**K**) Pie chart representing the extended ZGA genes according to their expression changes upon chemical inhibition of Nr5a2 (red - upregulated, blue - downregulated, gray - no change) and the total Nr5a2 CUT&Tag signal measured in their 8k bp upstream regions (dark - strong, mid shade - weak, light - no occupancy)

*De novo* motif analysis revealed that Nr5a2 motifs were enriched in 76% of Nr5a2 CUT&Tag peaks, whereas Esrrb motifs were enriched in 52% of Esrrb CUT&Tag peaks (Fig. 3C). The motifs that emerged from the detected peaks contained (A/G)(A/G)T upstream of the consensus sequences (Fig. 3D), which was also detected in motif 1 in the *SINE B1/Alu 5YR* upstream of ZGA genes (Fig. 1A, Fig. S1F). Indeed, we found that 70% of Nr5a2 peaks and 54% of Esrrb peaks overlapped with *SINE B1* (Fig. 3E and 3F), suggesting that *SINE B1* retrotransposons are major targets for Nr5a2 and Esrrb.

We examined the distance from the transcription start site (TSSs) of ZGA genes to the nearest Nr5a2 peak. We found that Nr5a2 peaks were substantially closer to ZGA genes (median: 14,059 bp) than to non-ZGA genes (median: 96,213 bp) (Fig. 3G). In addition, Nr5a2 peaks were much closer to the TSSs of ZGA genes that were downregulated vs. upregulated in Nr5a2 knockdown 2-cell embryos (Fig. 3G). We then analyzed the correlation between gene expression changes during ZGA and Nr5a2 occupancy of upstream regulatory regions. Nr5a2 occupancy was conservatively defined as the sum of Nr5a2 CUT&Tag signal over Nr5a2 motifs in the upstream 8 kb region of genes (Fig. S3F). We found that the gene expression changes between G2-phase zygotes and 2-cell stages was significantly higher for genes with Nr5a2 occupancy and stronger gene expression changes correlated with increased Nr5a2 occupancy (Fig. 3H and S3G). We further examined the correlation between gene expression changes in Nr5a2 knockdown and SR1848-treated embryos and Nr5a2 occupancy. Genes with Nr5a2 occupancy were significantly less expressed upon Nr5a2 perturbation compared to unoccupied genes. Stronger loss of gene expression also correlated with increased Nr5a2 occupancy (Fig. 3I, 3J, S3H and S3I). In a similar analysis, we found that ∼50% (921/1809) of the major ZGA genes downregulated by SR1848 treatment showed some Nr5a2 CUT&Tag signal in their extended promoter regions, whereas most of the unchanged or upregulated major ZGA genes had no Nr5a2 signal on their corresponding regulatory sequences (Fig 3K). Based on these data, we propose that Nr5a2 binding close to ZGA genes promotes their transcriptional activation at ZGA. However, it cannot be excluded that distant binding of Nr5a2 also contributes to ZGA regulation.

### Nr5a2 and Esrrb target cell-type specific enhancers

To identify the *cis*-regulatory elements (cREs) that are bound by Nr5a2 and Esrrb in 2-cell embryos, we examined published ATAC-seq and ChIP-seq (histone modification) data (*38, 39*) and found that these transcription factors bound to open chromatin (Fig. 4A). We classified enhancer-like signatures (ELS) as regions with high H3K27ac and ATAC-seq and low H3K4me3 signals, vs. promoter-like signatures (PLS) with high H3K4me3 and ATAC-seq and low H3K27ac signals (Fig. S4A and S4B). Nr5a2-binding was enriched at distal enhancer-like signatures (dELS) (Fig. 4B), whereas Esrrb-binding was detected at ELS, PLS and other regions (Fig. 4B). These findings suggest that Nr5a2 and Esrrb target common and distinct loci, and that Nr5a2 preferentially binds enhancers in 2-cell embryos.

**Figure 4.**
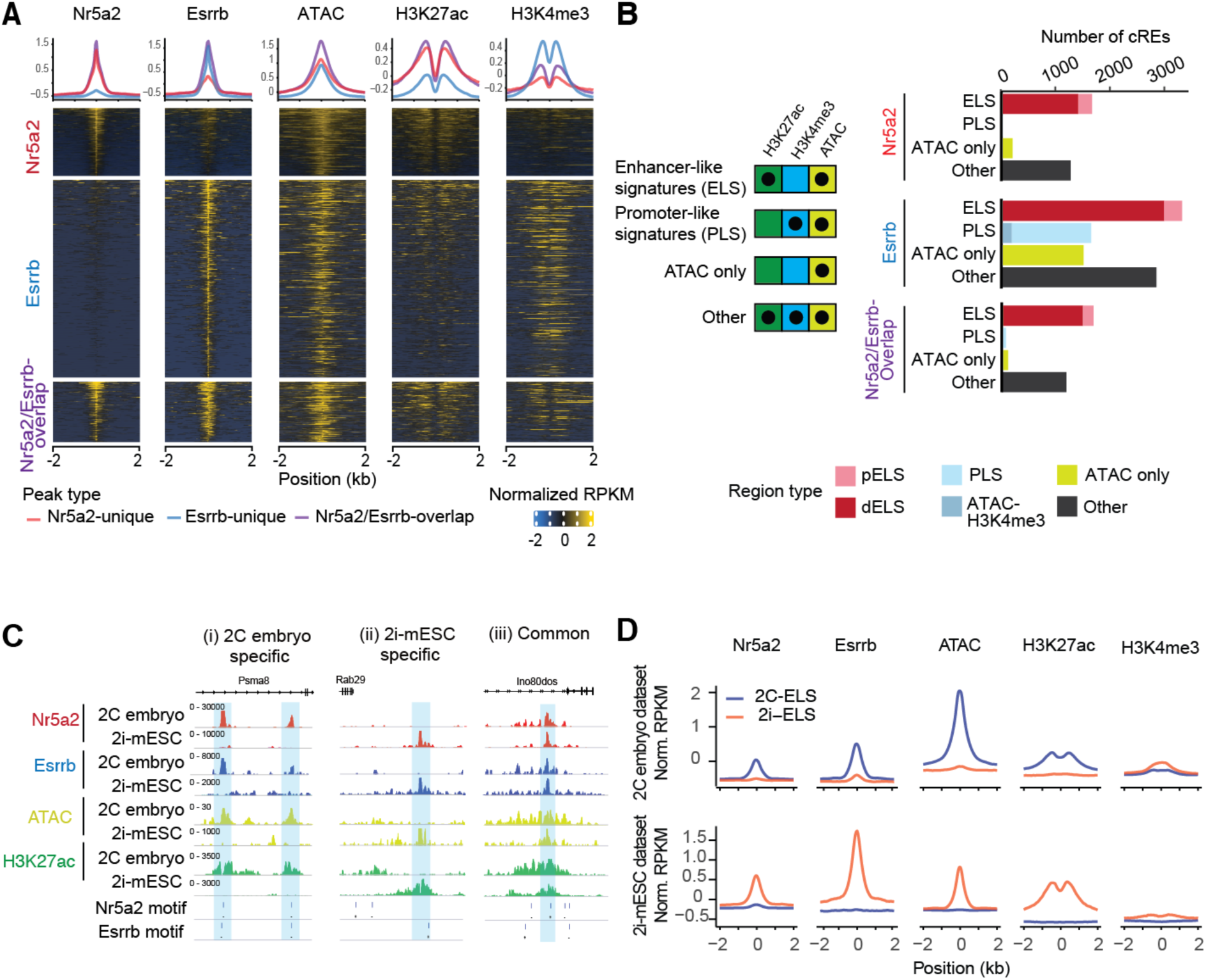
Cell-type specific distribution of Nr5a2 and Esrrb. (**A**) Line plots (above) and heatmaps (below) of Nr5a2, Esrrb, ATAC-seq, H3K27ac and H3K4me3 enrichments in 2-cell embryos (*Z*-score normalized RPKM value). Each row is classified by the Nr5a2 unique, Esrrb-unique and Nr5a2/Esrrb-overlap region. (**B**) Classification of cREs bound by Nr5a2 and Esrrb by epigenetic signatures. We defined 4 major groups: enhancer-like signatures (ELS), promoter-like signatures (PLS), ATAC only and other. Number of cREs bound by Nr5a2 and Esrrb are shown as bar graphs. The criteria of proximity are described in the method section. pELS: Proximal ELS. dELS: distal ELS. (**C**) Representative IGV snapshots show the enrichment of Nr5a2 (red), Esrrb (blue), ATAC-seq (yellow) and H3K27ac (green) in 2-cell embryo and 2i-mESC. Nr5a2 and Esrrb peaks are highlighted as blue. (**D**) Average profiles of Nr5a2, Esrrb, ATAC-seq, H3K27ac and H3K4me3 in 2C-ELS and 2i-ELS. The signal in a ± 2 kb window flanking the peak center is shown. Blue and orange lines indicate peaks on 2C-ELS and 2i-ELS, respectively.

Esrrb, and to a lesser extent Nr5a2, are expressed in ES cells (*26, 40*). To determine if these transcription factors are recruited to distinct cREs in totipotent 2-cell embryos vs. pluripotent ES cells, we examined published data (*40, 41*). Intriguingly, we found Nr5a2 and Esrrb binding to prominent ATAC-seq and H3K27ac peaks (ELS) that were (i) in 2-cell embryos only, (ii) in ES cells only, or (iii) in both 2-cell embryos and ES cells (Fig. 4C). We identified 9,099 2-cell-embryo-specific ELS (2C-ELS), 6,460 2i-mESC-specific ELS (2i-ELS) and 110 common ELS sites (Fig. S4C). Aggregation plot analysis showed that both Nr5a2 and Esrrb were specifically bound to each cell-type specific ELS with H3K27ac enrichment and chromatin accessibility (Fig. 4D, S4D and S4E). These data suggest that Nr5a2 and Esrrb may regulate gene-regulatory networks by defining cell-type specific enhancers, and that they bind to distinct cREs in pluripotent versus totipotent cells.

### Nr5a2 directly promotes chromatin accessibility

To test whether Nr5a2 and Esrrb are required for chromatin accessibility in 2-cell embryos, we developed a microscopy-based approach to quantify open chromatin in single cells, which we termed ChARM (Chromatin Accessibility Revealed by Microscopy) (Fig. 5A). Similar to ATAC-see (*42*), ChARM uses Tn5-mediated insertion of adaptor DNA into accessible chromatin but uses hybridization chain reaction (HCR) to amplify signals of adaptor DNA. This approach generated quantifiable spot-like patterns rather than the diffusive signals observed by ATAC-see (Fig. S5A and S5B). As a proof of concept, we examined Bromodomain 4 (Brd4)-dependent open chromatin. Two-cell embryos treated with the Brd4 inhibitor JQ-1 showed reduced H3K27 acetylation, as expected (Fig. S5C and S5D), and also significantly reduced ChARM signal (Fig. S5C and S5E).

**Figure 5.**
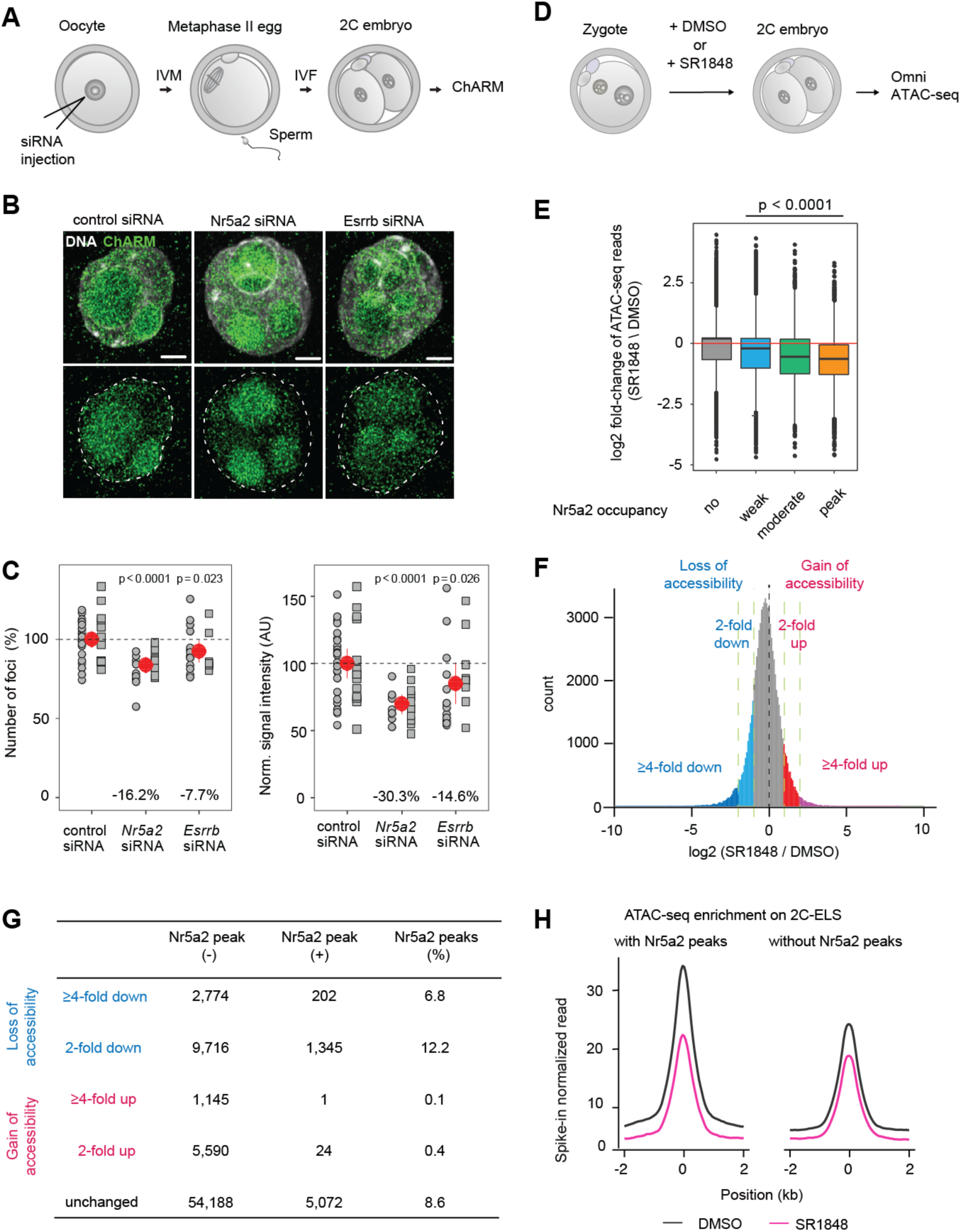
Nr5a2 regulates chromatin accessibility during ZGA. (**A**) Schematic illustration of ChARM. (**B**) Representative images of ChARM (green) in 2-cell embryos. DNA labeled with DAPI is shown in gray. Scale bars are 5 μm. (**C**) Scatterplot shows the relative percentage of the number of ChARM foci per nucleus and normalized signal intensity. Experimental replicates are shown with different bullet styles. Red dots and bars indicate the mean and 95% confidence interval of the experimental condition with all replicates merged. Number of nuclei analyzed in the replicates: control: n = 24, 13; *Nr5a2* knockdown: n = 9, 12; *Esrrb* knockdown: n = 14, 9. (**D**) Schematic illustration of Omni ATAC-seq in 2-cell embryos treated by DMSO or SR1848. (**E**) Boxplot of relative changes in ATAC-seq signal upon SR1848 treatment at Nr5a2 motifs with no (N = 42,158), weak (N = 26,712), moderate (N = 4,935) Nr5a2 signal or Nr5a2 signal strong enough for peak-calling (N = 5,029). Bonferroni corrected p-values of pairwise Mann-Whitney U tests against genes with no Nr5a2 occupancy are shown. (**F**) Histogram showing loss and gain of accessibility regions. (**G**) Summary of each chromatin accessibility regions with or without Nr5a2 peaks. (**H**) The average Omni ATAC-seq enrichment with or without Nr5a2 peak at 2C-ELS.

To test whether Nr5a2 and Esrrb promote chromatin accessibility, we performed knockdown in oocytes and analyzed 2-cell embryos by ChARM at 26 hpf. Chromatin accessibility was reduced in Nr5a2- and Esrrb-siRNA embryos compared to controls, suggesting that each transcription factor contributes to chromatin accessibility (Fig. 5B and 5C). Similar results were obtained for 2-cell embryos treated with SR1848 (Fig. S5F and S5G). Together, these findings imply that these transcription factors promote chromatin accessibility.

To determine whether Nr5a2 functions as a pioneer factor, we tested whether Nr5a2 is required for chromatin opening at sites where it is bound. SR1848 was used to inhibit Nr5a2 in 2-cell embryos and obtain sufficient cell numbers to perform Omni Assay for Transposase-Accessible Chromatin using sequencing (Omni ATAC-seq) (*43*) (Fig. 5D). SR1848 treatment resulted in a significant loss of chromatin accessibility at most Nr5a2 motifs with Nr5a2 occupancy and the magnitude of loss increased with occupancy (Fig. 5E). Motif analysis identified the Nr5a2 motif in loss of accessibility regions (Fig. 5F, S5H and S5I), which were also enriched for Nr5a2 peaks (Fig. 5G). The majority of Nr5a2 peaks were enriched in unchanged regions (Fig. 5G), implying that Nr5a2 promotes chromatin accessibility on a subset of its bound regions.

We next addressed whether Nr5a2-binding promotes chromatin accessibility on 2C-ELS. SR1848 treatment reduced chromatin accessibility more at 2C-ELS with Nr5a2 peaks than those without Nr5a2 peaks (Fig. 5H). These results suggest that Nr5a2 binding promotes opening of chromatin, which is a hallmark of pioneer factors.

### Nr5a2 and Esrrb bind nucleosomal DNA *in vitro*

Another hallmark of pioneer factors is the ability to target their (partial) motif on nucleosomal DNA (*23*). To test whether Nr5a2 and Esrrb possess this ability, we purified full-length and DNA-binding domains (DBDs) of mouse Nr5a2 and Esrrb, and mouse histones as recombinant proteins (Fig. S6A-D). Mass photometry showed that Nr5a2 forms a monomer, whereas Esrrb forms a dimer or multimer in solution (Fig. S6E).

To examine the binding specificity to naked DNA, we performed fluorescence polarization (FP) measurements and electrophoretic mobility shift analysis (EMSA). Since Esrrb showed promiscuous binding to DNA (Fig. S6F), we performed experiments in the presence of low concentrations of competitor DNA. Nr5a2 bound to its own motif and the Esrrb motif with comparable affinity (*K*_d_ of 5.62 ± 1 nM and 6.49 ± 0.35 nM, respectively). In contrast, Esrrb bound to its own motif with a higher affinity than the Nr5a2 motif (*K*_d_ of 8.56 ± 0.69 nM and 625.53 ± 133 nM, respectively) (Fig. 6A and S7G). The same motif specificity was observed by EMSA (Fig. S6H and S6I).

**Figure 6.**
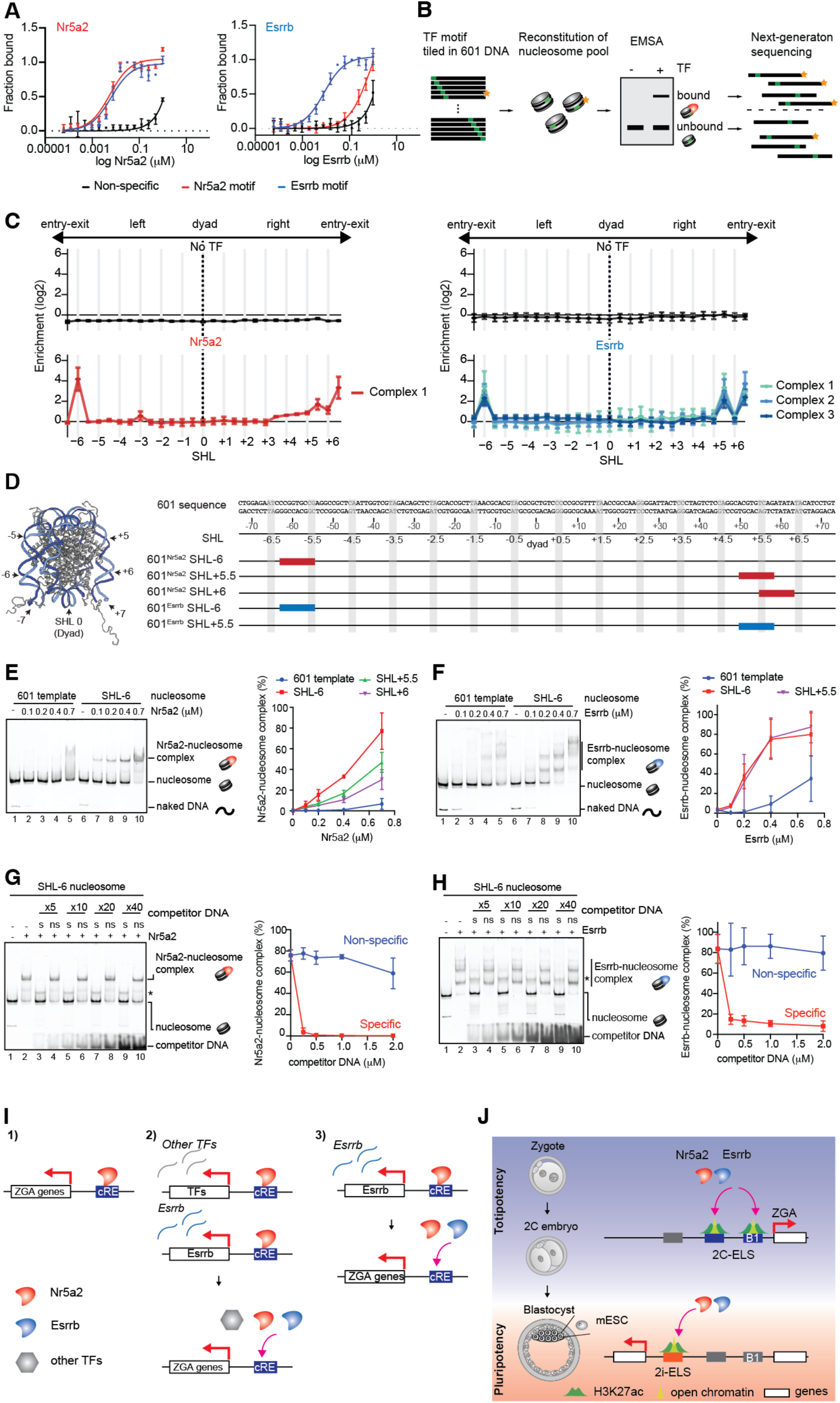
Nr5a2 and Esrrb specifically recognize nucleosomal target DNA. (**A**) DNA binding measured by fluorescence polarization for Nr5a2 (left) and Esrrb (right) using DNA with non-specific (black lines), Nr5a2 motif (red lines) and Esrrb motif (blue lines). The average values of three independent experiments are shown with the SD values. (**B**) Schematic illustration of SeEN-seq. Nucleosome libraries were reconstituted with TF motif (green) tiled in 601 DNA. TF-bound and unbound fractions were recovered and sequenced for revealing position-specific enrichments. Star indicates a sequence of specific enrichment as an example. (**C**) SeEN-seq enrichment profiles of Nr5a2 and Esrrb. The enrichments (log_2_) were plotted against each SHLs (from SHL −6.5 to SHL +6.5). The average values of two independent experiments are shown with the SD values. (**D**) Left panel shows a schematic of SHL positions on nucleosome (PDB ID: 1KX5). Right panel shows the location where Nr5a2 (red) or Esrrb (blue) motif is inserted on 601 DNA sequence. (**E and F**) Nucleosome binding assays with Nr5a2 (E) or Esrrb (F). Left panel shows representative data of EMSA with the 601 template nucleosome and SHL-6 nucleosome. Right panel shows graphical representation. The average values of three independent experiments are shown with the SD values. (**G and H**) Competition assay with Nr5a2 (G) and Esrrb (H). Nr5a2 or Esrrb (0.5 μM) was incubated with SHL-6 nucleosome containing their own motifs (50 nM) in the presence of 5-, 10-, 20- and 40-fold molar excess of specific competitor DNA (“s” lanes) or non-specific DNA (“ns” lanes). Asterisk indicates the competitor DNA-bound complex. Quantification of the results shown in right panel. The average values of three independent experiments are shown with the SD values. (**I and J**) A model on the Nr5a2-dependent ZGA regulation in mouse embryos.

To test whether Nr5a2 and Esrrb bind nucleosomal DNA, we performed SeEN-seq (Selected Engagement on Nucleosome sequencing), in which motifs are tiled throughout the Widom 601 nucleosome positioning sequence (Fig. 6B) (*44*). We prepared nucleosome libraries with 5 bp intervals of each motif (Fig. S7A, S7C, Table S7). EMSA showed a shift of Nr5a2 with the nucleosome library but not with the 601 “template” nucleosome lacking a motif, suggesting that the Nr5a2-nucleosome complex forms in a motif-dependent manner (Fig. S7B). Similarly, Esrrb showed specific band shifts with the nucleosome library (Fig. S7D). Transcription factor-bound and -unbound fractions were purified and sequenced. SeEN-seq revealed that Nr5a2 and Esrrb preferentially bound at the entry-exit sites on the nucleosome (Fig. 6C), reminiscent of how the pioneer factors Oct4-Sox2 and GATA3 binding to nucleosomal DNA (*44, 45*).

To investigate Nr5a2 and Esrrb binding at specific motif positions on nucleosomes, we selected high enrichment sites and reconstituted nucleosomes with motifs at superhelical locations (SHL) −6, +5.5 and +6 for Nr5a2; and −6 and +5.5 for Esrrb (Fig. 6D). Consistent with the SeEN-seq results, a bandshift was detected for Nr5a2 binding to nucleosomes containing the motif at SHL-6, SHL+5.5 and SHL+6 (Fig. 6E and S7F). Motif-specific binding was also detected for the Nr5a2 DBD. Esrrb full-length and DBD showed similar nucleosome binding efficiencies (Fig. 6E, S7G, and S7I). A stronger bandshift was detected for Esrrb binding to nucleosomes with motifs at SHL-6 and SHL+5.5 than for 601 template (Fig. 6E and S7G).

To test the specificity of the transcription factor-nucleosome interactions, we performed competition assays with naked DNA. Specific but not non-specific DNA outcompeted binding of Nr5a2 and Esrrb to nucleosomes (Fig. 6F and 6G). Thus, both Nr5a2 and Esrrb directly engage with their own motifs on nucleosomal DNA and preferentially target near the entry/exit sites of the nucleosome. Overall, our data provide evidence that Nr5a2 and Esrrb have properties consistent with pioneer factor activity *in vivo* and *in vitro*.

## Discussion

How ZGA is initiated in mammalian embryos is poorly understood. We show that the orphan nuclear receptor Nr5a2 is a pivotal pioneer factor that activates major ZGA genes in mouse embryos. Chemical inhibition suggests that 72% of ZGA genes are regulated by Nr5a2 and potentially other related orphan nuclear receptors. More than half of the ZGA genes are directly bound by Nr5a2, suggesting that it has a predominant role in regulating their expression. The remarkable genome-wide regulation of ZGA exceeds that of Nfya, which affects 15.1% of ZGA genes (*13*) and is comparable to Zelda in *Drosophila* (∼75% of ZGA genes) (*5*) and the combined action of three pluripotency factors in zebrafish (>75% of ZGA genes) (*8, 9*). The combination of genome-wide binding profiles, chromatin accessibility assays, and *in vitro* nucleosome binding provide evidence that Nr5a2 acts locally to promote chromatin opening and functions as a pioneer factor in mouse embryos. The classification of Nr5a2 and Esrrb as pioneer factors suggests that the mechanism of multiple pioneer factors triggering ZGA is evolutionarily conserved from fly to mouse and possibly human, even though the identity of the pioneer factors that trigger ZGA differs between species.

Our work shows that *SINE B1/Alu* retrotransposable elements are major targets of Nr5a2 in the genome. The binding of Nr5a2 to these elements in linear proximity to the TSS of ZGA genes correlates with their transcriptional regulation. Based on this, we consider a model in which Nr5a2 recruitment to *SINE B1/Alu* with enhancer-like properties activates the majority of ZGA genes. We cannot exclude that Nr5a2 recruitment outside canonical *SINE B1/Alu* (30% of the peaks) also contributes to ZGA regulation. However, these “isolated” Nr5a2 motifs show signatures of degenerate *SINE B1/Alu* elements, implying that most Nr5a2 motifs are derived from retrotransposon propagation (Fig. S3J).

In addition to Nr5a2 motifs, *SINE B1/Alu* contain motifs for other orphan nuclear receptor family members such as Esrrb and unrelated transcription factors. Indeed, both Nr5a2 and to a lesser extent Esrrb contribute to ZGA. Elucidating their regulatory network is complicated by the upregulation of both genes at ZGA. Our findings are consistent with several models (Fig. 6I): 1) Nr5a2 directly controls most ZGA genes, including other orphan receptors (e.g. *Esrrb*), 2) Nr5a2 controls expression of transcription factors including orphan nuclear receptors and these control most ZGA genes, and 3) Nr5a2 controls transcription of orphan nuclear receptors and the proteins co-operate in controlling most ZGA genes. Combinatorial approaches will be required to distinguish between these models.

An intriguing finding is that the same transcription factors are important for totipotency and later for pluripotency during mammalian development. Crucially, Nr5a2 and Esrrb target distinct enhancer-like sequences in 2-cell embryos and 2i-ES cells. The open question therefore remains of how pioneer factors selectively establish cell-type specific enhancers. *SINE B1/Alu* are transiently transcribed and accessible in 2-cell embryos but not in ESCs (*39, 46*). Whether *SINE B1/Alu* transcription facilitates access for pioneer and other transcription factors to define 2C-ELS remains to be determined. Since *SINE B1/Alu* contains several transcription factor motifs, we propose that cooperative transcription factor binding including Nr5a2, Esrrb and potentially other transcription factors establishes 2-cell embryo-specific active enhancers (Fig. 6J). Our findings imply that maternally provided Nr5a2 protein initiates a cascade of transcription factor bindings that lead to the transcriptional waves at the start of life.

## Acknowledgements

We would like to thank A. Mohanan, A. Lalic, B. Kunkel and E. Krstevska-Vulic for technical assistance. We thank all members of the K.T. laboratory including B. J. Dequeker, E. E. Chatzidaki, R. J. Ashburn, S. Feng and H. Marvanova for assistance with oocyte isolation. JQ-1 was a kind gift from J. Zuber (IMP). We thank R. S. Grand (Schübeler lab, FMI) for advice on SeEN-seq analysis. We thank R. H. Kim for sequencing single embryo samples at the NGS facility in the Department of Totipotency, MPIB. Illumina sequencing of the developmental transcriptome analysis was performed by the NGS facility at Vienna BioCenter Core Facilities (VBCF). We thank M. Novatchkova (IMP) for RNA-seq analysis of MZT data. We would like to thank J.-M. Peters and Life Science Editors for critical reading of the manuscript.

## Funding

L’Oréal Austria Fellowship for Women in Science (JG)

Austrian Science Fund (FWF) DK Chromosome Dynamics grant W1238-B20 (KT, JG)

JSPS Overseas Research Fellowship (WK)

European Research Council grant ERC-CoG-818556 TotipotentZygotChrom (KT)

Human Frontier Science program RGP0057-2018 (KT)

Austrian Academy of Sciences (KT)

Max Planck Society (KT)

## Author contributions

JG collected MZT samples for developmental RNA-Seq. JG, AV, LCR and LGH performed oocyte microinjections and collected samples for FISH and RNA sequencing analyses. JG performed Esrrb and Nr5a2 immunofluorescence assays. JG carried out embryo developmental competence assays. IG designed and carried out the ZGA-FISH experiments. IG and PK analysed single embryo RNA-seq data. WK and SL established the method of CUT&Tag. WK performed CUT&Tag and Omni ATAC-seq using mouse oocytes and embryos. SR, IG and PK analyzed CUT&Tag data. IG and JG performed ChARM and analysed data. SR and IG analysed Omni ATAC-seq data. WK, MK and MZ performed protein purification, SeEN-seq and biochemical analyses. SR analyzed SeEN-seq data. KT conceived the project and supervised the work. WK, IG, JG, SR, and KT planned the project, designed the experiments, and wrote the manuscript. All authors discussed the results and commented on the manuscript.

## Data and materials availability

Requests for plasmids generated in this study should be directed to the corresponding author. All RNA sequencing data have been deposited to the Gene Expression Omnibus (GEO) and will be made available upon publication. All CUT&Tag and Omni ATAC-seq data have been deposited to the Gene Expression Omnibus (GEO) and will be made available upon publication. ATAC-seq and histone modification ChIP-seq in 2-cell embryo datasets were downloaded from GEO accession GSE66390 and GSE72784, respectively. For the dataset in mESCs, ChIP-seq data of Nr5a2, Esrrb, and H3K27ac were obtained from GSE92412. H3K4me3 ChIP-seq was obtained from GSE56312.

Any additional information required to reanalyze the data reported in this paper is available from the corresponding author upon request.

