## Supplementary data for "Zygotic genome activation by the totipotency pioneer factor Nr5a2"

**Materials and Methods**

Animals

The care and use of the mice were carried out in agreement with the authorizing committee according to the Austrian Animal Welfare law and the guidelines of the International Guiding Principles for Biomedical Research Involving Animals (CIOMS, the Council for International Organizations of Medical Sciences). Mice were kept at a daily cycle of 14-h light and 10-h dark with access to food ad libitum. Mice were bred in the IMBA animal facility or purchased from Jackson laboratories (strain 101043, B6129SF1/J). Wild-type mice of B6129F1 background were obtained from breeding C57BL/6J females and 129/Sv males. To obtain zygotes and 2-cell embryos, mice were mated with B6CBAF1 stud males or oocytes were in vitro fertilized with B6CBAF1 sperm. For developmental RNA-Seq experiments pure B6 mice (C57BL/6J females crossed to C57BL/6J males) or B6CASTF1 mice (C57BL/6J females crossed to CAST/EiJ males) were used.

Collection of mouse oocytes and early embryo

For oocyte collection, ovaries of 8- to 20-week-old female mice were isolated and punctured with hypodermic needles to release the GV oocytes. Oocytes were isolated from 12-32 females per experiment, pooled and split into the different experimental conditions. For isolation of zygote and 2-cell embryos, 3- to 6-week-old female mice were mated with males. To induce superovulation, females were injected with PMSG (5IU) followed by hCG injection 48 hours later. Zygotes and 2-cell embryos were obtained by puncturing the oviduct after 17 hours (5 hpf) and 45-46 hours (33-34 hpf) post hCG injection, respectively.

In vitro maturation and in vitro fertilization (IVM/IVF)

In vitro maturation and fertilization were performed as described (44) with some changes. Oocytes were cultured in microdrops under 5% O2, 5% CO2, 90% N2. Oocytes were isolated in M2 medium supplemented with dbcAMP (Sigma, 300 µM) and FBS (20%) and microinjected for siRNA knockdown (see below) and further cultured for 5 hours in M16 supplemented with dbcAMP. Oocytes were released from dbcAMP and cultured for 12 hours in M16, followed by in vitro fertilization (In Vitro Fert, Cooks; supplemented with GSH (Sigma)). Fertilized embryos were cultured in KSOM until fixation for RNA FISH or ChARM.

Microinjection of oocytes for siRNA knockdown and rescue of knockdown

For siRNA knockdown, isolated GV oocytes were microinjected with sets of two siRNAs against targets (20 µM each) or control (20 µM each (Dharmacon) or 40 µM total (IDT)). As a control of successful injection 150 ng/µl H2B-EGFP (for control) or 150 ng/µl H2B-rsEGFP (for target) was co-injected. Predesigned siRNAs were obtained from following vendors: Nr5a2

(ON TARGET plus siRNA, Dharmacon); Esrrb (DsiRNA, IDT) using the respective controls from the vendors.
Used siRNAs were: Nr5a2: J-047044-09-002, J-047044-10-002; Control: ON-TARGETplus Non-targeting Control siRNA #1, ON-TARGETplus Non-targeting Control siRNA #2; Esrrb: mm.Ri.Esrrb.13.1, mm.Ri.Esrrb.13.2; Control: Negative Control DsiRNA (51-01-14-04);
For the rescue experiment, isolated GV oocytes were microinjected either as described above for siRNA knockdown or with addition of 500 ng/µl Nr5a2 mRNA.

<u>Chemical treatment of mouse embryos</u>
3-6 weeks old B6129F1 female mice were superovulated by injection of PMSG (5IU) followed by hCG injection 48 hours later. After injection, females were transferred into stud cages with B6CBAF1 male mice. Zygote were isolated after 17 hours post hCG injection (5 hpf) and transferred into potassium-supplemented simplex-optimized medium (KSOM) containing either 0.1% DMSO, triptolide (Sigma, 2 µM), 1.2 µM JQ-1 (THP Medical Products, HY-13030) or 5-10 µM SR1848 (Sigma, SML1513) at 6 hpf. The embryos were cultured in Lumox dish (Sarstedt) without mineral oil under high humidity conditions (wet chamber) until 34 hpf.

<u>Embryonic development assay</u>
Embryos were cultured in microdrops of KSOM using lumox dishes (35 mm, Sarstedt) under 5% O2, 5% CO2, 90% N2. Embryonic development was monitored once every 24 hours using a stereomicroscope. For inhibitor treatment zygotes were isolated at 5 hours post fertilization and cultured in inhibitors or mock control from 6 hours post fertilization onwards. Due to the hydrophobic nature of the inhibitors, embryos were cultured in wet chambers without oil.

<u>Immunofluorescence</u>
After oocyte or embryo collection from B6129F1 mice, cells were fixed in 4% PFA diluted in PBS for 1h, before permeabilization in 0.5% Triton X-100/PBS (PBT) for 30 min. Blocking was carried out in 1% BSA (Sigma) in 0.5% PBT for 1h at room temperature. Cells were incubated overnight at 4°C with primary antibody (anti-Nr5a2, 1:100 (Origene, TA805074, Clone: OTI3H8, Lot: W001); anti-Esrrb, 1:100 (Perseus Proteomics, PP-H6705-00), anti-H3K27ac, 1:500 (Active Motif, 39133) diluted in 1% BSA (Sigma) in 0.5% PBT. After washing in blocking solution for at least 30 min, incubation with the secondary antibody (Alexa Fluor 568 goat anti-mouse IgG (H + L), dilution 1:500, Thermo Fisher Scientific #A-11031, Lot: 1736975; Alexa Fluor 488 goat anti-rabbit IgG (H + L), dilution 1:500, Thermo Fisher Scientific #A-11008, Lot: 645232) in 1% BSA (Sigma) in PBT was carried out for 1 h at room temperature. Another set of washing steps in 0.1% PBT (PBS + 0.1% Triton X-100) were followed by incubation with 1.25 µg/ml DAPI in PBT for 20 min. After washing in PBS, cells were mounted in Vectashield (Vector Labs) and imaged on a confocal microscope (LSM880, Zeiss, ZEN blue) using a 63×, 1.4NA oil objective.

<u>Low-input and single embryo RNA-seq</u>
Triplicates of small bulk samples of 10 embryos each or single embryos in 2-3 replicate collections were collected, the zona pellucida was removed with Acidic Tyrode's solution (Sigma) and briefly washed through 3 drops of 1x PBS. Embryos were lysed in 3 µl of 0.2% Triton X-100 and 2U RNaseOUT in PCR tubes as described previously (45) and kept on ice for >10 min and snap frozen in liquid nitrogen until further sample preparation. For developmental RNA-Seq, MII eggs, zygotes (G1 and G2-phase) and 2-cell embryos were collected in triplicates of small bulk format (10 eggs/embryos) at 14, 19.5, 26 and 46 hours post hCG injection respectively (corresponding to 7.5, 14 and 34 hours post fertilization for zygotes and embryos). Sequencing libraries were prepared using SmartSeq v4 kit (Takara Bio) and Nextera XT (Illumina) according to the manufacturer's protocol. Sequencing was performed in the VCBF Next Generation Sequencing facility using paired-end 50 bp sequencing on HiSeq2500 (Illumina, HiSeqv4).
For single embryo sequencing, mock and inhibitor treated cells or control and knock-down samples were collected as single embryos at 34 hours post fertilization (56 hours post hCG

injection). Inhibitor treatment and knock-down were performed as described above. Sequencing libraries were prepared using the SmartSeq2 protocol described in52. Preamplification PCR was performed using Q5 High-Fidelity Master mix (NEB) and 12 cycles. Further library preparation was performed using Nextera XT (Illumina) according to the manufacturer's protocol. The libraries were sequenced using NextSeq 500 for paired end sequencing using High Output Kit v2.5 (400M reads).

FISH probe design

To probe gene expression during ZGA, corecommon major ZGA genes were chosen as targets for nascent and mature ZGA-FISH, respectively. Genes that have at least 1500 bps of obligatory intronic or exonic sequences - that is the sequences in question were intronic or exonic in all annotated isoforms encoded by the locus - were identified as potential targets. grouped into three non-overlapping (ZGAlow, ZGAmod and ZGAhigh) categories to robubstly based on their transcript abundance in 2-cell embryos to design nascent ZGA probes. For robust detection of *de novo* ZGA transcripts, three non-overlapping sets of potential target genes (N = 81, 79, 81)were selected intofrom these three categories and 50-50 unique probes with 22-28 nucleotide complementary regions to target were designed as described previously (46). Probes were extended with barcodes at both ends and an incomplete T7 promoter at their 3′ end (see Table S5) and were synthesised as a ssDNA oligo pool (Genscript). Category specific, fluorescently labeled probe arrays were produced by PCR amplification (ZGA#1low: 5′ catatacgctcgtcgggact + 5′ atctTAATAcgactcactatagggtattttacg, ZGA#2Amod: 5′ ttcaaggatgatccgcgctt + 5′ acatTAATAcgactcactataggcatgatctag, ZGA#3high: 5′ aattaggctcggtcgcccta + 5′ aaacTAATAcgactcactatagggttgacggac; nucleotides in capital represent the missing part of the T7 promoter to be completed during the PCR). Resulting dsDNA was bead-purified (in-house magnetic beads) and transcribed in vitro to RNA with high-yield T7 kit (Thermo or NEB). RNA was bead-purified as above and its molar concentration was determined. Using equimolar fluorescently labeled oligos (5′ [Atto565]acgctcgtcgggact, 5′ [Atto647]ggatgatccgcgctt and 5′ [Atto532]cggtcgcccta) RNA was reverse transcribed into fluorescent ssDNA. RNA template was removed by O/N digestion with RNaseH and ssDNA was bead-purified. Quality of probe arrays was assessed on denaturing 10% PAGE (8M urea) and by absorbance measurements at 260 nm at dye-absorption maxima.

ZGA - Fluorescent In Situ Hybridization (ZGA-FISH)

Embryos at the appropriate developmental time were fixed with 4% PFA in PBS for 20 minutes. Following wash-out and quenching of PFA with Tris-HCl for 10 min, the embryos were washed twice in PBT and dehydrated by a passage through a series of ethanol (25%, 50%, 75% and 2x 100%, diluted with PBT). Subsequently they were stored at -20 °C until use. Prior to hybridization, the embryos were rehydrated through a series of dilution ethanol (75%, 50%, 25% diluted in PBT), then washed twice in PBT. Prehybridization was done in 1x pre-HYBEC (2x SSC, pH 7.0, 1 mM EDTA, 15% ethylene carbonate and 0.1% Triton X-100) supplemented with 50 µg/ml heparin at 42 °C for 60-90 minutes by placing the embryos into a humid chamber in a water bath. 10 minutes before the end of prehybridization the hybridization mixture was prepared: 1x pre-HYBEC, 50 µg/ml heparin + 40-40 nM ZGA#1low and ZGA#2 probe arrays (~10-10 pM/unique probe; ZGA#3high was reserved as backup and not used routinely due to dye incompatibility between the co-injection marker GFP and Atto532 in our imaging setup) and was heated to 65°C for 5 minutes to denature intra- and intermolecular dimers in the pools. Preheated hybridization mixture was applied to the embryos and hybridization was carried out for 6-7 h or O/N at 42°C similarly as prehybridization.

At the end of hybridization, embryos were washed in prewarmed 1x pre-HYBEC three times for a total of 1 h at 42 °C , followed by two-three short washes in PBT or 1x SSCT (1xSSC + 0.1% Triton X-100) at room temperature (RT). DNA was counterstained with 1.25 µg/ml DAPI either in the second PBT wash (then this would last for 20 min) or during the equilibration with the Abberior Liquid Mounting medium (20 min).

2-cell embryos in which chromatin accessibility was also imaged were first tagmented as described in the next section, then used for ZGA-FISH after stopping the tagmentation and subsequently were recovered for developing the ChARM signal. To facilitate recovery, coverslips were not sealed onto the slides. They were gently lifted off and embryos were collected into 1x SSCT. Probes from previous experiments were removed by dehybridization: embryos were incubated 3 x 15 min in DEHYB buffer (0.1x SSCT, 1 mM EDTA, 30 % ethylene carbonate, 0.1% Triton X-100) at 55-60°C. Dehybridized embryos were washed 2 x 5 minutes in 1x SSCT at RT, followed by the prehybridization step of the subsequent experiment.

Chromatin Accessibility Revealed by Microscopy

Rehydrated 2-cell embryos for ChARM analysis were washed in 1×TMP buffer (10 mM Tris-HCl, pH 8.3, 5 mM MgCl2, 4% PEG 8000) after the second PBT wash (see FISH section above). 20 µM Tn5 preloaded with p7#1 (5'GTCTCGTGGGCTCGGCTGTCCCTGTCCCGAGTAATCACCGTCTCCGCCTCAGATGT GTATAAGAGACAG) and p5#1 (5' TCGTCGGCAGCGTCTCCACGCTATAGCCTG CGATCGAGGACGGCAGATGTGTATAAGAGACAG) sci-ATACseq adapter oligos38 for ChARM or Atto 594 labelled ATAC-see adapter oligos24 or Atto 565 labelled p7#1 and p5#1 oligos for ATAC-see in 1×TMP buffer was added to the embryos and those were incubated at 37°C for 30-60 minutes. From this step on, the embryos were handled in a humid chamber. Tagmentation was stopped and DNA bound Tn5 was removed from chromatin by 3×5 minutes wash in STOP buffer (10 mM EDTA, 0.05% SDS in 1×SSC) at room temperature. Embryos were then washed 2×5 minutes in 1x SSCT (1×SSC + 0.1 % Triton X-100) at RT. After this step, KD and SR1848-treated embryos were first assayed with ZGA-FISH. For ATAC-see, embryos were mounted after this step (see below). For ChARM, the embryos - fresh or recovered from ZGA-FISH - were pre-hybridized in 1×preHYBEC buffer (2×SSC, 1 mM EDTA, 15 % ethylene carbonate, 50 µg/ml heparin and 0.1% Triton X-100) at 37°C for 30 minutes. 2-2 µM split-initiator oligonucleotides (p7#1-B5-1: 5' CTCACTCCCAATCTCTATAA AGGGACAGCCGAGCCCACGAGA, p7#1-B5-2: 5' GAGGCGGAGACGGTGATTACTCGGGAACTACCCTACAAATCCAAT, p5#1-B5-1:5'CTCACTCCCAATCTCTATAACGTGGAGACGCTGCCGACGA, p5#1-B5-2: 5' TGCCGTCCTCGATCGCAGGCTATAACTACCCTACAAATCCAAT) were pre-warmed to 37°C for 1-2 minutes and were applied to the pre-hybridized embryos. Hybridization was carried out overnight at 37°C. Next morning excess initiator molecules were washed away by 3×20 minutes in 1×preHYBEC without heparin. During the last wash, fluorescently labeled B5 H1 and H2 HCR hairpins21 were prepared by heating up to 80 ℃ and cooling to 25 ℃ at 0.5 ℃/s in a thermocycler. Embryos were washed for 2×5 minutes in 5×SSCT (5xSSC + 0.1 % Triton X-100). Prepared H1 and H2 hairpins were diluted to 3-3 µM in 5×SSCT and were applied to the embryos at RT for 6-7 hours in the dark. After the HCR, embryos were washed for 4-5×15 minutes in 5×SSCT at room temperature in the dark. DAPI staining was done either during one of these washes (typically the 3rd) or during the mounting using 1.25 µg/ml DAPI in 5×SSCT (wash staining) or in Abberior Liquid Mounting Medium (mounting staining). Embryos were transferred into Abberior Liquid Mounting Medium for 20 minutes for equilibration.

In case of the JQ-1 treatment, 2-cell embryos were stained with anti-H3K27ac antibody to test the efficiency of HAT inhibition. Briefly, embryos were recovered after imaging, washed in PBT then blocked in BLOCK (1% BSA and 1:50 Dernhardt's solution (Sigma) in PBT). Anti-H3K27ac (Active motif #39133) diluted 1:500 in BLOCK were applied to the samples overnight at 4 ℃. Embryos were washed 4×15 minutes in PBT at RT. AlexaFluor647 conjugated anti-rabbit Fab fragments (Jackson ImmunoResearch) diluted 1:1000 in BLOCK were applied to samples for 2 h at RT in the dark. Embryos were washed 4×15 minutes in PBT at RT and were mounted for a second round of imaging.

Images were restored by deconvolution in SVI Huygens Professional. Whole nuclei and cytoplasm were segmented in Z-stacks that cover the entire nuclear volume using a custom-built macro in ImageJ. ChARM foci were detected in these segmented images by a custom spot detector plug-in. Data of the detected spots were analyzed in R using RStudio. Spot intensity normalization was done per nucleus by fitting a series of Gaussian functions to the signal intensity distribution of the corresponding cytoplasmic spots. The smallest μ (typically half of the μ in the series, an indication of a good fit) was used to normalize the nuclear spot intensities. Nucleolar foci were filtered out based on overlap with minimal DNA intensity regions (below the 4th percentile of intranuclear DAPI signal intensity distribution) within the nuclei, typical of nucleolar bodies. Total number of the detected nuclear foci and total normalized signal intensity were compared to control. Multiple replicates of the same experiments were combined by further normalizing the data to the mean of the corresponding control.

<u>Imaging and image analysis</u>

After the equilibration, the embryos are mounted on slides with pre-printed wells and imaged on a Zeiss LSM 880/980 point scanning confocal microscope using a 63x 1.4 NA oil immersion objective. Detector settings were adjusted for sequential scans of Atto565 (561 nm ex, 575-610 nm em), GFP (to monitor the expression of the co-injection marker, 488 nm ex, 495-545 nm em) and Atto647 (639 nm ex 650-720 nm em) together and DAPI (405 nm ex, 420-470 nm em). 4x oversampled stacks (63 nm lateral and 250 nm axial resolution) were collected according to modified Nyquist criteria, which were subsequently deconvoluted by Huygens Professional (SVI).

Deconvolved images were segmented in 3D into nuclei (full nuclear volumes) and surrounding cytoplasm using a custom ImageJ macro in FIJI/ImageJ. DNA and H2B-GFP signal intensity were measured in the nuclear compartment. Spots representing the FISH signal were detected using a custom ImageJ plugin (47, 48). Collected data was analyzed in R using RStudio (www.rstudio.com). Briefly, after rejecting samples based on undetectable H2B-GFP co-injection marker (if applicable; typically 0-2 nuclei per sample), spots detected in the cytoplasm - that may represent leaking pre-mRNA from the nuclei in case of the nascent ZGA-FISH or tagmented mtDNA in case of ChARM - were used as a basis of compensating for imaging artifacts (e.g. scattering) that result in signal intensity differences. On an individual stack-by-stack basis, the signal intensity distribution of cytoplasmic spots was fitted with a series (K) of Gaussian functions using the mixtools package as described previously (46). A good fit was obtained if the smallest and the subsequent μ of the Gaussians had an approximate ratio of 1:2, typically at K=2 or 3. Using the smallest μ, all raw intensity values (cytoplasmic and nuclear) collected from the given stack of images were normalized. The normalized cytoplasmic distribution showed a high fraction (>80%) of single-μ spots. As all the transcription units are expected to be present within each nuclei in the same amounts - that is we did not expect any large scale aneuploidy and the embryos are expected to be in a similar developmental stage - the normalized values were summed up within individual nuclei and were the basis of the analysis to derive total nuclear signal (or copy number). To merge multiple replicates of the same experiment, data was further normalized to the mean of the corresponding control to compensate for day-to-day variances in the observed numbers.

Data was visualized by ggplot2. Statistical comparisons were carried out by two-tailed pairwise nonparametric Mann-Whitney U tests with $\alpha = 0.05$ with Bonferroni correction when multiple testing was performed on the experiment (and not replicate) level. Significance of individual replicates can be judged by the span of the 95% confidence intervals - whether they cross the 100% reference line - plotted as error-bars. Due to the uncertainties of in vitro embryonic development, sample sizes were not determined prior to the experiment.

<u>Preparation of pA-Tn5 or Tn5 adaptor complex</u>

The 3×Flag-pA-Tn5-Fl plasmid (Addgene plasmid # 124601) was purchased from Addgene. Recombinant pA-Tn5 was purified according to published protocol (32). In-house Tn5 was provided from the molecular biology service in Vienna Bio Center. The concentration of pA-Tn5 was determined by SDS-PAGE with recombinant Tn5 as the standard protein. To prepare the pA-Tn5 or Tn5 adaptor complex, 10 μl of 5.5 μM pA-Tn5 or Tn5 was mixed with 1.6 μl of a 100 μM equimolar mixture of Tn5MEDS-A and Tn5MEDS-B oligonucleotides (32). The mixture was incubated at room temperature for 1-2 h before dilution for CUT&Tag or Omni ATAC-seq.

CUT&Tag on mouse GV oocytes and 2-cell embryos

CUT&Tag was performed as described previously (32) with a few modifications. We performed H3K4me3 CUT&Tag on GV-stage oocytes and Nr5a2 CUT&Tag on 2-cell embryos with normal high salt condition (300 mM NaCl). Esrrb CUT&Tag on 2-cell embryos was performed with physiological salt condition (150 mM NaCl) due to its low binding affinity to the genome. Isolated GV oocytes and 2-cell embryos with intact zona pellucida were incubated with ice-cold extraction buffer (25 mM HEPES-NaOH (pH 7.4), 50 mM NaCl, 3 mM MgCl2, 300 mM sucrose and 0.5% Triton X-100) on ice for 10 min and 7 min, respectively. Cells were washed three times through an ice-cold extraction buffer without Triton X-100. Pre-extracted cells were further lightly fixed by DPBS with 0.1% formaldehyde for 2 min at room temperature. To stop the reaction, cells were washed through 5 drops of M2 buffer. Polar bodies in 2-cell embryos were carefully removed by a micromanipulator combined with a piezo drive unit (PMM-150FU, Prime Tech). Cells were briefly washed with 9 µl of antibody buffer (20 mM HEPES-NaOH (pH7.5), 150 mM NaCl, 0.5 mM Spermidine, 0.1% BSA, 2 mM EDTA and 1x Protease inhibitor cocktail (Roche)) in a 10 µl well of a microplate (NuncTM MicrowellTM Minitrays). We placed 10-20 cells/well. Cells were incubated with 9 µl of antibody buffer with primary antibody (1:100 dilution, anti-H3K4me3: #04-745 Merck, anti-Nr5a2: ABE2867, Merck, anti-Esrrb: PP-H6705-00, Perseus Proteomics) overnight at 4°C. To remove primary antibody, cells were washed three times for each 20 min through 9 µl of antibody buffer and further incubated with secondary antibody (1:100 dilution, Guinea pig anti-rabbit (Heavy & Light chain) antibody (ABIN101961, Antibodies-Online) for anti-rabbit primary antibody; rabbit anti-mouse IgG H&L (ab46540, abcam) for anti-mouse primary antibody) for 1.5 h at room temperature. Cells were washed three times for each 20 min through 9 µl of Wash buffer (20 mM HEPES-NaOH (pH7.5), 150 mM NaCl, 0.5 mM Spermidine and 1x Protease inhibitor cocktail (Roche)) to remove unbound antibodies. From pA-Tn5 binding step, we used Wash-300 buffer (20 mM HEPES-NaOH (pH7.5), 300 mM NaCl, 0.5 mM Spermidine and 1x Protease inhibitor cocktail (Roche)) for high salt conditions; and Wash buffer for physiological conditions. pA-Tn5 adaptor complex was prepared by 1:250 dilution with Wash-300 buffer/Wash buffer. Cells were incubated with diluted pA-Tn5 for 1.5 h at room temperature and washed three times for each 20 min through 9 µl of Wash-300 buffer/Wash buffer to remove unbound pA-Tn5. Cells were transferred into 1.5 ml of DNA LoBind Tube (Eppendorf) with 200 µl of Tagmentation buffer (10 mM MgCl2 in Wash-300/Wash buffer) and incubated at 37°C for 1h. To stop tagementation, 10 µl of 0.5 M EDTA, 2 µl of 10% SDS and 2 µl of 20 mg/ml Proteinase K (Roche) was added to sample, and sample were incubated at 50°C for 2 h for deproteinization and then further incubated at 65°C for 16 h for reverse crosslinking. After a quick spin to remove liquid from the cap, the sample was brought to total 300 µl by adding 0.1×TE buffer. The DNA was extracted by phenol-chloroform and was precipitated with ethanol and 1.5 µl of 20 mg/ml glycogen. The precipitation mix was incubated at -80 °C for 30 min or overnight, followed by centrifugation at 21,130 x g at 4°C for 30 min. The pellet was washed with 200 µl of 70% EtOH, followed by centrifugation at 21,130×g at 4°C for 20 min. The pellet was resuspended in 20 µl of 0.1×TE buffer, and 1 µl of 1 mg/ml RNaseA was added to the sample. After RNase-treatment at 37°C for 10 min, the DNA was amplified 50 µl of reaction with 1×Q5 High-Fidelity Master mix (NEB) and 0.25 µM barcoded primers using the following PCR program: 72 °C for 5 min; 98°C for 30 sec; 15 cycles of 98°C for 10 sec and 63°C for 30 sec; final extension at 72 °C for 1 min and hold at 4°C. After PCR reaction, libraries were purified by 1.1×AMPure XP magnetic beads (Beckman Coulter) and were eluted in 20 µl of RNase-free water. Purified DNA libraries were analyzed by BioAnalyzer (Agilent Technologies) and were sequenced on a NextSeq 500 with paired-end 75-base pair (bp) reads (Illumina). See Table S6.

Omni ATAC-seq on 2-cell embryos

Zona pellucida was removed by treatment with acidic Tyrode. Omni ATAC-seq was performed as described previously (40) with a few modifications. Embryos were transferred into 1.5 ml of DNA LoBind Tube (Eppendorf) with 200 µl of PBS and were spun at 500×g at 4°C for 5 min to remove supernatant. 50 µl of ATAC-Resuspension buffer (RSB) (10 mM Tris-HCl (pH 7.4), 10 mM NaCl, 3 mM MgCl2) containing 0.1% NP-40, 0.1% Tween-20 and 0.01% Digitonin was added into tube and incubated for 3 min on ice. After incubation, 200 µl of ATAC-RSB containing 0.1% Tween-20 was further added into the tube to wash out lysis. Nuclei were spun at 500×g at 4°C for 10 min to remove supernatant. Nuclei were incubated with 50 µl of transposition mixture (10 mM Tris-HCl (pH 7.6), 5 mM MgCl2, 10% Dimethyl formamide, 33% PBS, 0.1 % Tween-20, 0.01% Digitonin and 95 nM Tn5-adaptor complex) at 37°C for 30 min with 1000 rpm mixing. To stop tagementation, 10 µl of 0.5 M EDTA, 2 µl of 10% SDS, 2 µl of 20 mg/ml Proteinase K (Roche) and 150 µl of 300 mM NaCl was added to sample, and sample were incubated at 50°C for 1 h. After a quick spin to remove liquid from the cap, the sample was brought to a total 300 µl by adding 0.1×TE buffer. The DNA was purified by phenol-chloroform and followed by ethanol precipitation with 1.5 µl of 20 mg/ml glycogen and 2 µl of Drosophila spike-in DNA(49). The precipitation mix was incubated at -80 °C for 30 min or overnight, followed by centrifugation at 21,130×g at 4°C for 30 min. The pellet was washed with 200 µl of 70% EtOH, followed by centrifugation at 21,130×g at 4°C for 20 min. The pellet was resuspended in 20 µl of 0.1×TE buffer, and 1 µl of 1 mg/ml RNaseA was added to the sample. After RNase-treatment at 37°C for 10 min, the DNA was amplified 50 µl of reaction with 1×NEBNext HF 2×PCR Master mix (NEB) and 0.25 µM barcoded primers using the following PCR program: 72 °C for 5 min; 98°C for 30 sec; 15 cycles of 98°C for 10 sec, 63°C for 30 sec and 72 °C for 1 min; final extension at 72 °C for 5 min and hold at 4°C. After PCR reaction, libraries were purified by 1.1×AMPure XP magnetic beads (Beckman Coulter) and were eluted in 20 µl of RNase-free water. Purified DNA libraries were analyzed by BioAnalyzer (Agilent Technologies) and were sequenced on a NextSeq 500 with paired-end 75bp reads (Illumina). See Table S6.

Protein expression and purification

The DNA fragment encoding Mus musculus Nr5a2 (560 aa, UniProtKB: P45448) was ligated into NdeI-BamHI sites of pET-15b vector. His6-tagged Nr5a2 was expressed in E.Coli BL21 (DE3) codon plus RIL (Agilent Technologies). The cells were cultivated at 30 °C until an optical density of 0.6 (OD595), and His6-tagged Nr5a2 expression was induced with 0.25 mM IPTG at 16°C for 16-18 h. The cells were resuspended in buffer 1 (20 mM Tris-HCl (pH 7.5), 300 mM NaCl, 15 mM imidazole, 5% glycerol, 2 mM CHAPS and 1×protease inhibitor cocktail (Roche)) and were disrupted by sonication. The cell debris was removed by centrifugation. The supernatant was gently mixed with Ni-NTA agarose beads (Qiagen) at 4°C for 30 min. The beads were packed into an Econo-column (Bio-Rad) and were washed with 50 column volumes (CVs) with buffer 1. Protein bound to the Ni-NTA agarose beads was eluted by a linear gradient of 15-500 mM imidazole. Fractions containing His6-Nr5a2 were dialyzed against buffer 2 (20 mM Tris-HCl (pH 7.5), 400 mM NaCl, 5% glycerol, 2 mM CHAPS and 1 mM DTT) for further purification. After dialysis, the sample was diluted with buffer 3 (20 mM Tris-HCl (pH 7.5), 5% glycerol, 2 mM CHAPS and 1 mM DTT) to reach 150 mM NaCl concentration. The diluted sample was immediately loaded onto the cation exchange column (MonoS 5/50 GL (Cytiva) or SP Sepharose Fast Flow (Cytiva)). The column was washed with 20 CVs of buffer 4 (20 mM Tris-HCl (pH 7.5), 150 mM NaCl, 5% glycerol, 2 mM CHAPS and 1 mM DTT), and the sample was eluted by a linear gradient of 150-900 mM NaCl. The purified sample was dialyzed against buffer 5 (20 mM Tris-HCl (pH 7.5), 400 mM NaCl, 5% glycerol and 1 mM DTT). Aliquots of purified protein were frozen at -70°C.

The DNA fragment encoding Mus musculus Esrrb (433aa, UniProtKB: Q61539) was ligated into NdeI-BamHI sites of pET-15b vector with a N-terminal His6-TEV tag. Essrb was purified by the same method as Nr5a2 purification except for His6-tag removal with TEV protease.

Nr5a2 DBD (100-208 aa) and Esrrb DBD (97-194 aa) were cloned into pET-28a vector. The sequence encoding SUMO-His6-tag was added at the C-terminus of Nr5a2 DBD and Esrrb

DBD. Both Nr5a2 DBD and Esrrb DBD were purified by the same method as Nr5a2 purification except for SUMO-His6-tag removal with PreScission protease.
The DNA fragments encoding mouse H2A (Hist1h2ab), H2B (Hist1h2bc), H3.3 and H4 were synthesized by polymerase chain reaction (PCR). The amplified DNA fragments encoding mouse histones were ligated into the NdeI-BamHI site in the pET-15b vector. Mouse histones H2A, H2B, H3.3 and H4 were expressed and purified according to published protocols (50). For the reconstitution of histone octamer, lyophilized histones were equally mixed in denaturing buffer (20 mM Tris-HCl (pH7.5), 1 mM EDTA, 7 M guanidine hydrochloride, and 20 mM 2-mercaptoethanol). The reconstituted histone octamer was purified by size exclusion chromatography (Superdex 200 16/600, GE Healthcare). The purified histone octamer was concentrated with an Amicon Ultra centrifugal filter unit (Millipore) and stored at -70°C.

Mass Photometry
Imaging chambers for Mass Photometry were prepared as described previously (51). 20 µl of 25 nM of recombinant preparations of Nr5a2 or Esrrb in a buffer (20 mM Tris-HCl (pH 7.5), 120 mM NaCl, 1 mM $MgCl_2$, 10 µM $ZnCl_2$, 1 mM DTT) were applied into the flow cell and single-molecule binding events to the glass surface were imaged at room temperature (24˚C) on Mass Photometer (Refeyn One MP). 250-1000 binding events were recorded per sample. Obtained histograms were fitted with a Gaussian curve and the mean peak was determined using DiscoverMP software (Refeyn). Obtained distribution was calibrated with a standard curve as described previously (52). Each sample was measured at least 3 times independently.

DNA preparations
The DNA fragments for nucleosome reconstitution were amplified by PCR and further purified by polyacrylamide gel (6%) electrophoresis using a Prep Cell apparatus (Bio-Rad). The eluted DNA was concentrated with an Amicon Ultra centrifugal filter unit (Millipore).

Purification of nucleosomes
The DNA and mouse histone octamer were mixed in a 1:1.6-2.0 molar ratio in 2M KCl. The nucleosomes were reconstituted by salt dialysis method and were further purified by polyacrylamide gel (6%) electrophoresis using a Prep Cell apparatus (Bio-Rad). The nucleosomes were concentrated with an Amicon Ultra centrifugal filter unit (Millipore).

SeEN-seq assay
SeEN-seq assay was performed as described previously with a few modifications (41). Briefly, Nr5a2 motif (JASPAR ID: MA0505.1, TCAAGGCCA) or Esrrb motif (JASPAR ID: MA0141.1, TCAAGGTCA) was tiled 5 bp interval across the entire Widom 601 DNA sequence (Lowary and Widom, 1998). The individual DNA fragments were amplified by PCR and digested by EcoRV. Generated 153bp 601 DNAs containing Nr5a2 or Esrrb motif were purified by polyacrylamide gel (6%) electrophoresis using a Prep Cell apparatus (Bio-Rad). The purified DNA library was spiked with 601 DNA template (29:1 molar ratio; pool:601). Nucleosome pools were reconstituted and purified as described above. The pooled nucleosomes (0.1 µM) were incubated with Nr5a2 (0.4 µM) or Esrrb (0.4 µM) at room temperature for 30 min in a reaction buffer (20 mM Tris-HCl (pH7.5), 120 mM NaCl, 1 mM $MgCl_2$, 10 µM $ZnCl_2$, 1 mM DTT, 100 µg/ml BSA). After the incubation, the samples were separated by non-denaturing polyacrylamide gel (4.5%) electrophoresis. The bands were visualized by SYBR Gold staining (Invitrogen) using Gel DocTM XR+ system (Bio-Rad). Unbound and bound fractions were excised using GVM30 UV transilluminator (Syngene). DNA libraries were prepared as described previously (41). Purified DNA libraries were analyzed by BioAnalyzer (Agilent Technologies) and were sequenced on a NextSeq 500 with paired-end 150 bp reads (Illumina). See Table S7.

Fluorescence polarization assay

5 nM 5'-Alexa488-labeled 24 bp dsDNA containing Nr5a2 consensus motif (GAGAGAGTCAAGGCCATGGCTCACT), Esrrb (GAGAGAGTCAAGGTCATGGCTCACT) or non-specific sequence (shuffled, GAGAGAG GACTACGACTGGCTCACT) was used as a tracer. Serial 2-fold dilutions of Esrrb or Nr5a2 (0-2 μM) were mixed with 5 nM tracer in a reaction buffer (20 mM Tris-HCl (pH 7.5), 120 mM NaCl, 1 mM MgCl2, 100 μg/ml BSA, 10 μM ZnCl2, 1 mM DTT, 6.25 ng/μl Poly(dI-dC) (Thermo Fisher #20148E)) in a total volume of 20 ul and incubated for 30 min at RT in 384-well microplate (Grainer Bio-One, #781076). Changes in fluorescent polarization were measured by a PHERAstar FS microplate reader (BMG Labtech) equipped with a polarization filter. The polarization units were converted to fraction bound and plotted against increasing concentration of Nr5a2 and Esrrb in log10 scale. Prism (GraphPad) was used for curve fitting (one binding site model) and subsequently the apparent dissociation constant in the presence of competitor values (Kd_apparent) were obtained. All measurements were performed in triplicates.

Electrophoretic mobility shift assay (EMSA)

For EMSA with naked DNA, 50 nM of non-labeled or 5'-Alexa488-labeled 24 bp dsDNA containing Nr5a2 consensus motif (GAGAGAGTCAAGGCCATGGCTCACT), Esrrb consensus motif (GAGAGAGTCAAGGTCATGGCTCACT) or non-specific sequence (shuffled, GAGAGAGGACTACGACTGGCTCACT) was used a labeled probe. 0, 0.1, 0.2, 0.4 and 0.7 uM of full-length or isolated DNA-binding domain (DBD) of Nr5a2 or Esrrb were incubated with respective labeled probe in total reaction volume of 10 μl (20 mM Tris-HCl (pH 7.5), 120 mM NaCl, 1 mM MgCl2, 100 μg/ml BSA, 10 μM ZnCl2, 1 mM DTT) with or without 6.25 ng/ul Poly(dI-dC) (Thermo Fisher #20148E)) for 30 min in RT. Following addition of 2.5 ul of 30% sucrose, the samples were separated by non-denaturing polyacrylamide gel (10%). In the absence of competitor DNA, the bands were visualized by SYBR Gold staining (Invitrogen) using Gel DocTM XR+ system (Bio-Rad). In the presence of competitor DNA, Alexa488 fluorescence in-gel fluorescence was detected using BioRad Imager. The data was analyzed using ImageLab (Biorad) and plotted in Prism (GraphPad).

For KD measurement of DBD, 0.5 nM of 5'-Cy5, 3'-Cy5-labeled 24 bp dsDNA was incubated with serial dilutions of DBD (0-4 μM) in the same buffer conditions in the presence of 6.25 ng/ul Poly(dI-dC). Cy5 in-gel fluorescence was detected using BioRad Imager. The data from at least 3 replicates was analyzed using ImageLab (Bio-Rad) and plotted in Prism (GraphPad). Error bars show mean and standard deviation.

For EMSA with nucleosome, the nucleosomes (100 nM) were incubated with Nr5a2 (0, 0.1, 0.2, 0.4 and 0.7 μM) or Esrrb (0, 0.1, 0.2, 0.4 and 0.7 μM) at room temperature for 30 min in a reaction buffer (20 mM Tris-HCl (pH7.5), 120 mM NaCl, 1 mM MgCl2, 10 μM ZnCl2, 1 mM DTT, 100 μg/ml BSA). After the incubation, the samples were separated by non-denaturing polyacrylamide gel (4.5%). The bands were visualized by SYBR Gold staining (Invitrogen) using Gel DocTM XR+ system (Bio-Rad).

Statistical analysis

All statistical analysis was performed in R using non-parametric pairwise two-tailed Mann Whitney U tests as normal distribution of samples could not always be established. For rejecting the null-hypothesis, $\alpha=0.05$ was used with Bonferroni correction when multiple testing was performed. Prior testing, replicates were pooled. However, mean and 95% confidence intervals of individual replicates are shown relative to the mean of the corresponding control. Due to the uncertainties of in vitro embryonic development, sample sizes were not determined prior to the experiment.

Transcriptome analysis

Transcriptomes of small batches of MII eggs, zygotes and G2 stage 2-cell-embryos were analyzed in triplicates by pseudo-aligning paired-end sequencing reads to Mus musculus cDNA template (mm10) using Kallisto (53) without bootstrapping (kallisto quant). Data was imported to R by the tximport package (54) and analyzed on the gene level. Samples were

compared pairwise to the previous and subsequent developmental stages (if applicable) by DESeq2 (55). Sample relationships were determined by PCA after vst transformation. Genes were clustered with k means based on their expression similarities.
To specify major ZGA genes, genes with a minimum 4-fold increase (FDR=0.01) in expression between G2 zygote and G2 2-cell embryonic states were selected from the two datasets. The resulting list was further filtered for genuine robust expression change by keeping only those that had higher than 0.5 TPM abundance or higher than 50 TPM abundance in G2 zygotes and G2 2-cell embryos, respectively. This resulted in 2232 and 1261 major ZGA genes in the B6 x CAST and B6 x B6 datasets, respectively. The overlapping set of 985 genes were used for motif search and to design ZGA-FISH probes.
Single embryo transcriptomes were analyzed by pseudo-aligning paired-end sequencing reads to Mus musculus cDNA template (mm10) supplemented with the transcripts of EGFP, rsEGFP (co-injection markers) and the intronic sequences of Nr5a2. Pseudo-alignment of paired-end reads was performed by Kallisto (53) with 100 bootstraps . Data was imported to R by the tximport package (54) on the transcript level. After initial quality assessments and removal of technical outliers, Nr5a2 knockdown samples and their controls were batch corrected by BatchCorrectedCounts from the CountClust package (56) as replicate based clustering of the data was observed (for chemically inhibited embryos, no such correction was necessary). Samples were then compared by DESeq2 (55). Fold-changes between control and treated (RNAi or chemical inhibition) specimens were estimated using apeglm. PCA analysis was carried out following vst transformation.
For Nr5a2 knockdown, the knockdown efficiency was assayed by comparing the summed Nr5a2 transcript abundances between control and knockdown embryos. The minimum value observed in a control embryo was used as a threshold to split the Nr5a2 knockdown sample into strong and weak/moderate KD categories. In case of the SR1848 treatment, all inhibitor-treated embryos show below threshold values for Nr5a2 expression. One control embryo was deemed as an outlier based on a very low number of reads (including minimal Nr5a2 abundance) and was excluded from downstream analysis. Differentially expressed transcripts (FDR=0.1 and 0.05 for Nr5a2 knockdown and chemically inhibited embryos, respectively) were cross referenced with the two developmental transcriptome datasets. Individual genes (Nr5a2 or Esrrb) and their (premature) transcripts were analyzed in R using ggplot2 and pairwise Mann Whitney U test for statistical comparisons.

Motif search and genome feature analysis

To find enriched motifs, motif search with GADEM (Li, 2009) was performed using 8kb sequences upstream of the TSS of the 985 common major ZGA genes. The enriched motifs (Fig. S1F) were subsequently localized in the 8kb upstream sequences of common, B6 x CAST and B6 x B6 specific major ZGA genes (2508 in total) and of 6900 non-ZGA genes by the matchPWM function of the Biostrings package to calculate mean motif occurrences and fraction of genes with motifs. During visual inspection of motif positions we identified repeating patterns of motif co-occurrences. Analyzing these patterns, centered around motif #5 and #6, we confirmed the presence of these patterns. The consensus sequence of the resulting assembly of motifs (supermotif) matches that of SINE B1 with 90.3% identity.

CUT&Tag, ChIP-seq, ATAC-seq data processing

Low quality reads were trimmed out using TrimGalore (version 0.6.2) (https://github.com/FelixKrueger/TrimGalore) using parameters: --quality 20 --length 20. Read mapping was done using Bowtie2 (version 2.3.5.1) (57) using parameters: -t -q -N 1 -L 25 -X 2000 --no-mixed --no-discordant. Reads from CUT&Tag and published ChIP-seq and CUT&Run libraries (33-35, 37, 38) were mapped to mm10 genome. The read from ATAC-seq in 2-cell embryos (36) were mapped to N-masked reference genome, which was prepared from SNPs information in mm10 genome of C57BL/6NJ and DBA/2J strains (Mouse Genome Project) using SNPsplit (version 0.3.2)'s SNPsplit_genome_preparation function (58). Unmapped reads, low mapping quality reads (Q<30) were removed. PCR duplicated reads were identified and discarded using the MarkDuplicates command from Picard (version

2.18.27) (http://broadinstitute.github.io/picard/) with parameters: VALIDATION_STRINGENCY=LENIENT. The RPKM values were calculated to represent read coverage using the bamCoverage function from deepTools (version 3.1.2)(59). The RPKM values were further transformed using Z-score normalization for the visualization.
Reads from Omni-ATAC-seq datasets were mapped to both mouse (mm10) Drosophila (dm6, spike-in) genomes. Unmapped, low mapping quality, and PCR duplicated reads were removed as described previously. Spike-in normalized read count on each base is calculated by dividing read counts on mm10 genome by total spike-in reads (10,000/total mapped spike-in reads).
Reads from both DMSO and SR1848 were pooled together and used for peak calling (see below) to identify common peak set between the two samples. Differentially accessible peaks were classified based on the fold-change of the mean spike-in normalized coverage within the peak between SR1848 and DMSO treatment.

Nr5a2 occupancy
To calculate the Nr5a2 occupancy of upstream regulatory regions per gene, we summed the raw Nr5a2 CUT&Tag signal that overlapped with Nr5a2 motifs in the 8k bp region upstream of the TSS. This summed value was then used to represent the Nr5a2 occupancy of the given locus and was used to test correlations with changes in gene expression during ZGA and upon Nr5a2 interference. For statistical testing, the genes were sorted into no (0 read), weak (1-2 RPM), moderate (3-5 RPM) and strong (> 6 RPM) Nr5a2 occupancy categories and the genes with no occupancy were treated as baseline. The same Nr5a2 motif positions accumulated no reads in the corresponding IgG control (Fig. S3F).

Peak calling and motif analysis
Peak calling for Nr5a2 and Esrrb CUT&Tag datasets were done using findPeak command from HOMER (version 4.10)(60) with '-style factor' parameter. Peak calling of ATAC-seq and ChIP-seq were done using MACS2's callpeak function (version 2.2.5) with default parameters (61).

The reference motifs of Nr5a2 and Esrrb from JASPAR database (accession number MA0505.1 and MA0141.1, respectively) were used for the identification of their motif locations in the reference mouse genome (mm10). Motif searching was done using FIMO from MEME Suite (version 5.0.4) (62). The identification of known motifs and de novo motif analysis for both differentially accessible ATAC peaks and transcription factor CUT&Tag peaks were done using findMotifsGenome command from HOMER (version 4.10) with the '-size given' parameter.

Enrichment analysis of peak overlap
The enrichment analysis was performed in order to estimate the relative occurrence between observed and random overlapping of two genomic features, such as peaks of transcription factor and repeat elements, or cREs and TF peaks. The number of query regions (e.g. TF peaks) that contain at least one target region (e.g. repeat elements) within 500 bp from their centers was observed. The same approach was applied to randomly distributed query regions. Enrichment of the overlap is represented as log2-ratio between observed and random overlap. The genomic locations of repeat elements used in this analysis were obtained from RepeatMasker (https://www.repeatmasker.org/species/mm.html). Other genomic regions-related analysis, such as determining distance between peaks and transcriptional start sites, was done in R using the GenomicRanges package (63).

Defining cis-regulatory elements
Cis-regulatory elements in 2-cell embryo and mESC were identified based on the definition proposed by the ENCODE project (64). Regions with open chromatin, as defined by ATAC-seq peaks, and enriched in H3K27ac, and H3K4me3 are classified as regions with enhancers-like and promoters-like signature (ELS and PLS), respectively. Accessible chromatin regions that overlapped with both histone modifications are classified as 'other' regions. The ELS and

PLS are further classified into proximal and distal regions. ELS regions within 2,000 bp, and PLS regions within 200 bp from the closest transcriptional start site (TSS) are considered as proximal regions. The overlap between peaks was done using Bedtools (version 2.29.2) (65). Centers of ATAC-seq peaks were used as a reference point for further analyses of cis-regulatory elements in 2-cell embryo and mESC.

SeEN-seq data processing

Reads from SeEN-seq library were mapped into Widom 601 sequences that contain Nr5a2 or Esrrb motifs in 5 bp-window sliding variants using Bowtie2 (version 2.3.5.1) with parameters: -t -q --very-sensitive --no-discordant --no-mixed. Reads with mapping quality lower than 20 were discarded. The number of reads aligned to each construct was counted by Samtools idxstat command (66).

The binding enrichment was represented as described previously (41). In brief, the read count of each construct was normalized by the library size of each fraction. The library-size normalized count of each SHL construct was then divided by the normalized count of Widom 601 template sequence (67). Enrichment scores of each motif position are calculated as a log2-transformed fold-change between bound and unbound fractions.

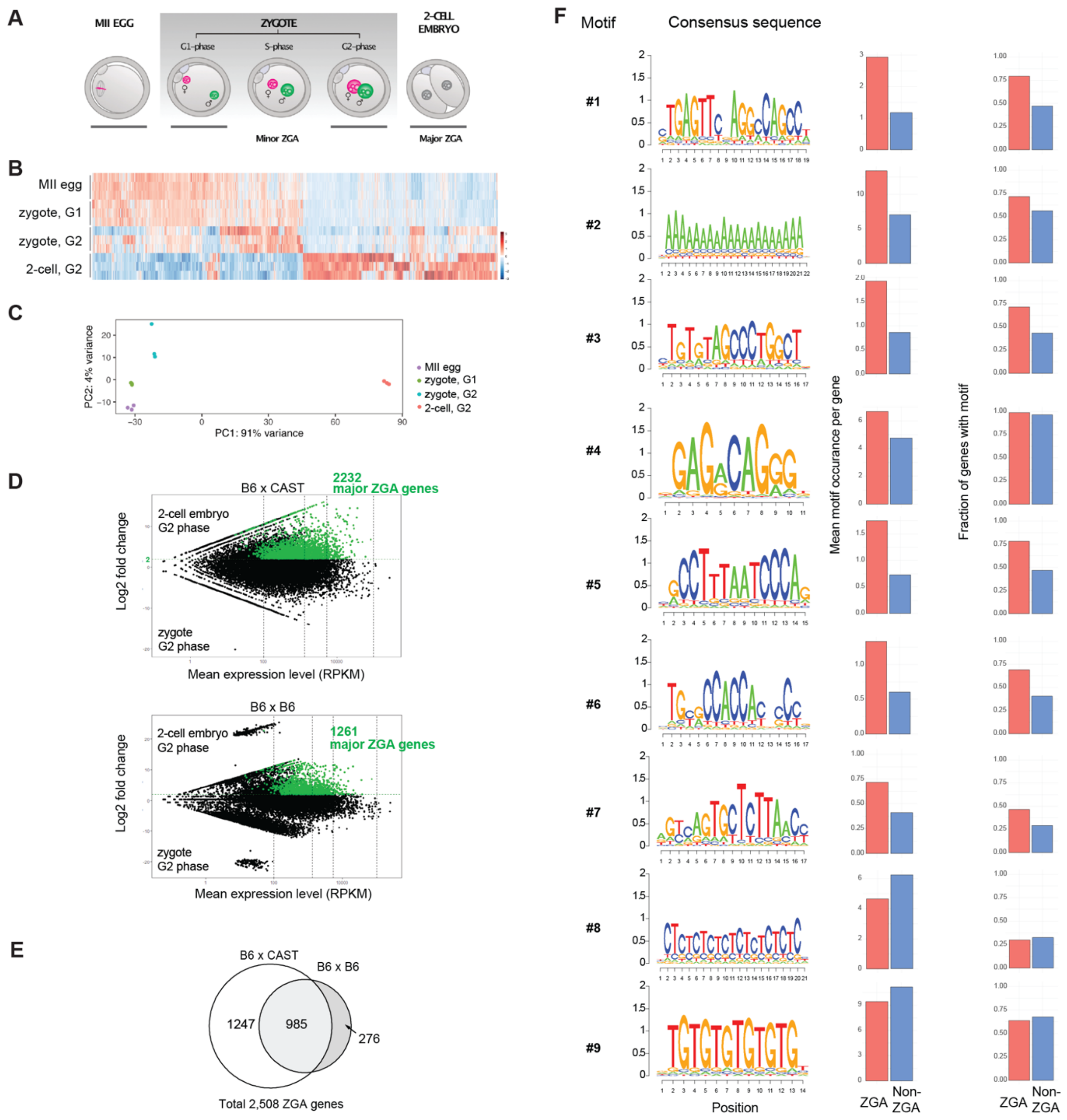


**Figure S1. Screening of potential transcription factors**

(**A**) Schematic depiction of embryonic stages in respect to minor and major ZGA. Collected stages for RNA-Seq are indicated by grey bars. (**B**) Comparison of developmental stages by cluster analysis of expressed genes. Stages were collected in triplicates. (**C**) Principal component analysis of the top 500 genes of the RNA-Seq data. (**D**) MA-plots of comparing gene expression between zygote G2- (bottom) and 2-cell embryo G2 phases (top) in B6CASTF1 hybrid (B6xCAST, Table S1) and C57BL/6J homozygous (B6xB6, TableS2) embryos. Green coloured dots indicate genes that met the selection criteria for ZGA. Numbers of ZGA genes indicated in the top right corners. (**E**) Euler diagram of overlapping ZGA genes between the two datasets. (**F**) Sequence logos of motif enriching in the 8 kb upstream regions from the TSS of the 985 common ZGA genes. Mean number of motif occurrence and fraction of genes that contain at least one motif in their 8kb upstream regions for all 2508 ZGA genes (red) and the 6900 non-ZGA genes (blue) used as control.

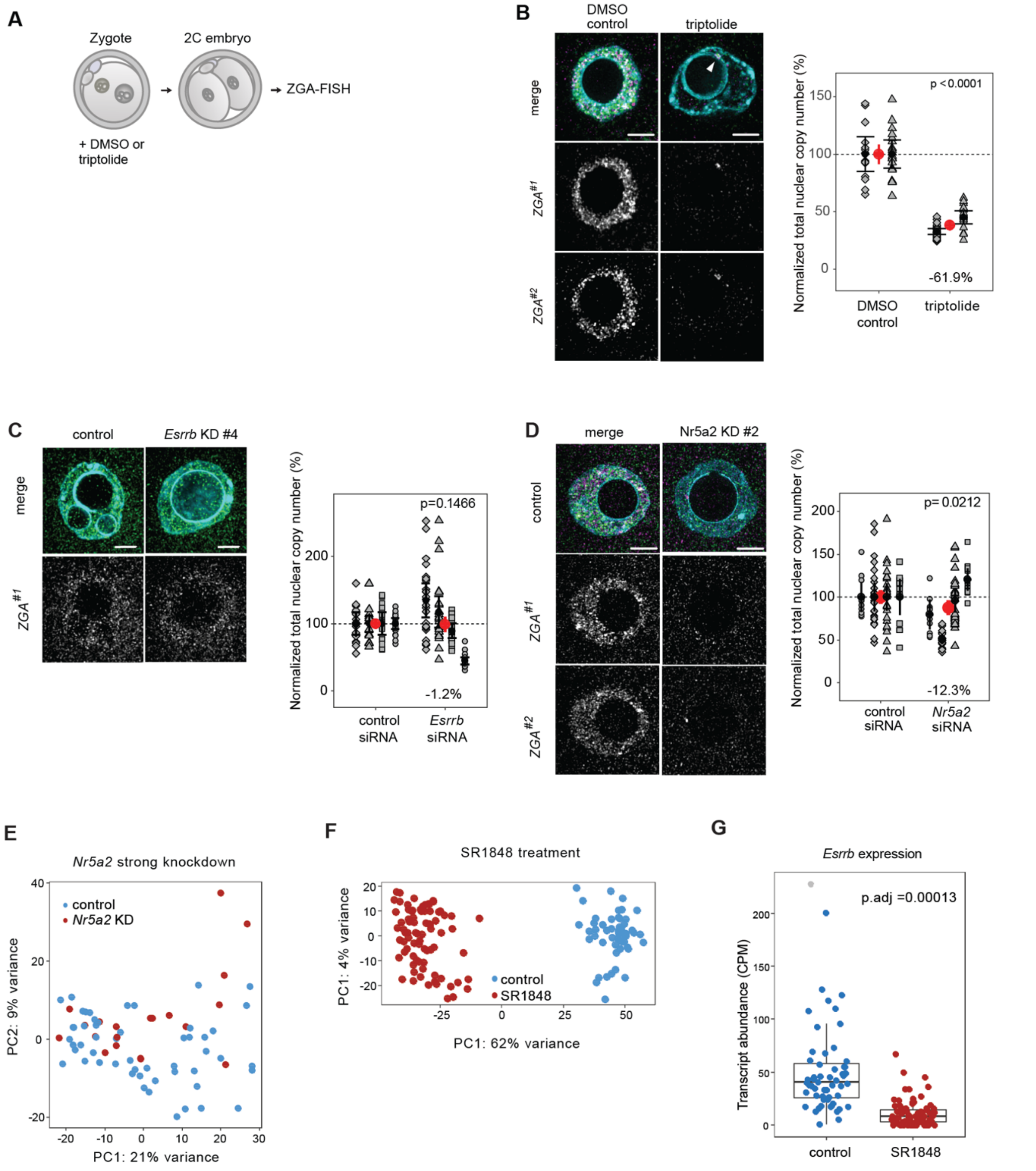


**Figure S2. Transcriptome analysis of *Nr5a2* knockdown and SR1848 treated 2-cell embryos**

(**A**) Schematic illustration of ZGA-FISH with DMSO control or triptolide treatment. (**B**) Representative images of nascent ZGA-FISH in nuclei of 2-cell embryos (34 hpf) treated with 2 µM triptolide. Probes against two sets of ZGA genes ($ZGA^{\#1}$ and $ZGA^{\#2}$) are shown in green and magenta, respectively, and DNA appears in cyan. Arrow indicates one of the few transcriptional foci resistant to triptolide treatment. Scale bars represent 5 µm. Quantifications of total nascent ZGA-FISH signal within nuclei of 2-cell embryos in two replicates are shown in the right panel. Each measurement per nucleus from replicate experiments is normalized to the mean of the corresponding control and plotted in different shapes. Mean and 95% confidence intervals of individual replicate experiments are indicated by black dot and error bars, mean and 95% confidence intervals of the pooled data are shown in red. P-value of pairwise nonparametric Mann Whitney U tests (α=0.05) between pooled experimental and control distributions is shown above, relative difference between the pooled experimental and control data is shown below the graphs. Sample sizes: control: n = 16, 19, triptolide: n = 27, 20 nuclei. (**C and D**) ZGA-FISH with knockdown 2-cell embryos (34 hpf). Representative images of ZGA-FISH (*ZGA*$^{\#1}$ - green, *ZGA*$^{\#2}$ - magenta) of the strongest effect showing *Esrrb* knockdown experiment (#4, circles in C) and *Nr5a2* knockdown experiment (#2, diamonds in D) assayed in 2-cell embryos. Scale bar is 5 μm. Quantification of total nascent ZGA-FISH signal within nuclei of *Esrrb* knockdown, *Nr5a2* knockdown and control 2-cell embryos in four replicates. P-value of pairwise Mann Whithney U test (α=0.05) is indicated in the top, the relative difference between the mean of the two populations is shown at the bottom of the graph. Scale bars represent 5 μm. Sample sizes in panel C are control: n = 17, 27, 17, 21; *Esrrb* knockdown: n = 25, 25, 18, 22 nuclei. Sample sizes in panel D are control: n = 12, 26, 16, 14; *Nr5a2* knockdown: n = 11, 16, 14, 14 nuclei. (**E and F**) Principal component analysis of strong Nr5a2 knockdown (E) and SR1848 treated 2-cell embryos (F) after DESeq2 analysis and vst transformation. (**G**) Abundance of *Esrrb* transcripts in 2-cell embryos treated with 10 μM SR1848.

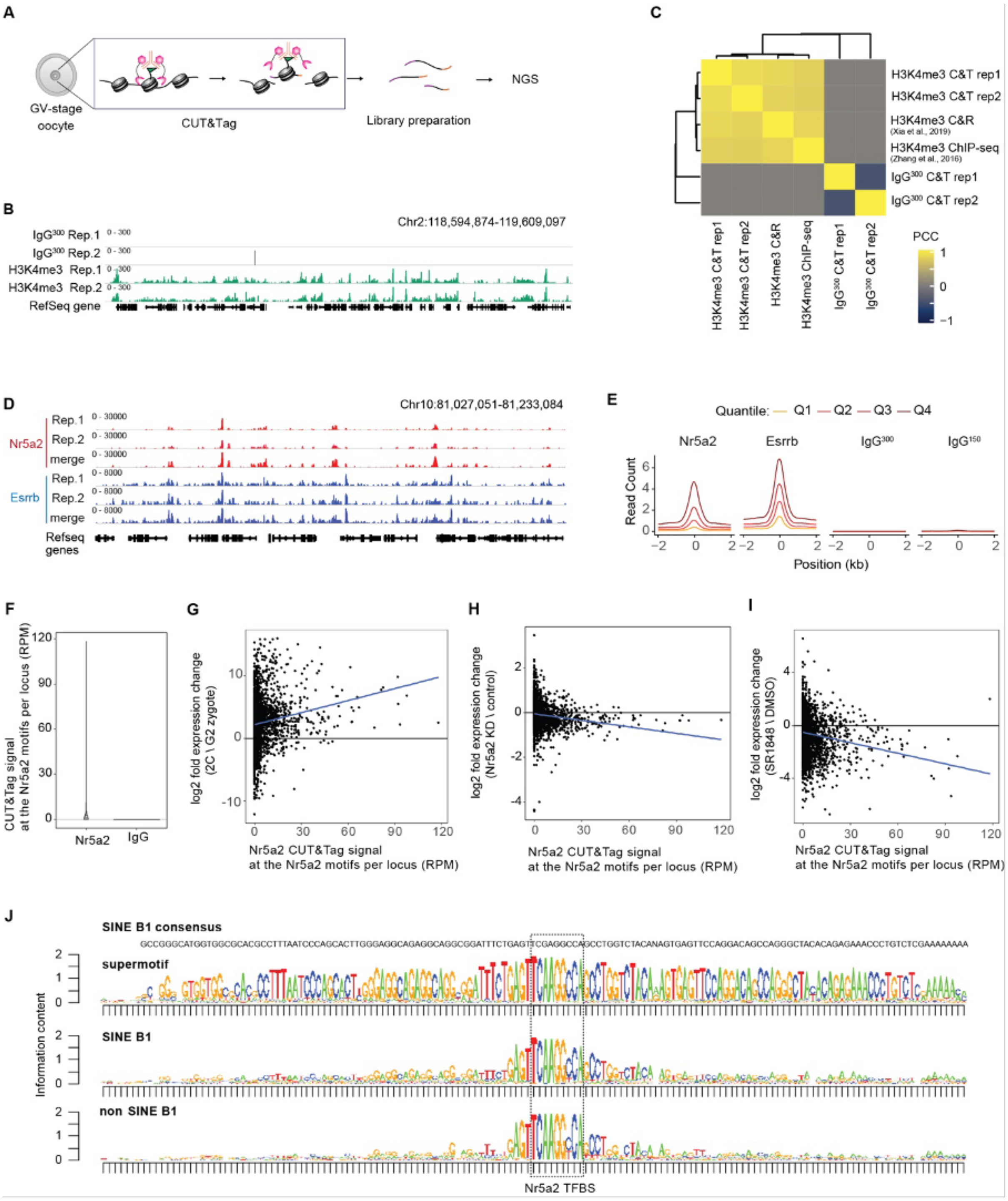


#### Figure S3 Validation of CUT&Tag

(**A**) Schematic illustration of CUT&Tag on mouse GV-stage oocytes. (**B**) Representative IGV snapshot shows the enrichment of IgG control (grey), H3K4me3 (green). (**C**) Correlation matrix (Pearson correlation) comparing between our H3K4me3 CUT&Tag (C&T) and public ChIP-seq (33) and CUT&Run (C&R) (34) dataset in mouse oocytes. (**D**) Representative IGV snapshot comparing between two replicates of Nr5a2 and Esrrb CUT&Tag on 2-cell embryos. (**E**) Aggregation plot shows read counts of each region. The data are colored according to the quantile range in the ATAC-seq data. (**F**) Violin plot of summed Nr5a2 and IgG CUT&Tag reads at Nr5a2 motifs in the upstream 8 kbp region of genes. (**G-I**) Correlation analysis between Nr5a2 occupancy of Nr5a2 motifs in the 8kbp upstream regions of genes and their expression change (F) between G2 zygotes and 2-cell embryos, (G) in Nr5a2 KD and (H) in SR1848-treated 2-cell embryos. (**J**) Sequence logos around the Nr5a2 motifs (indicated by the dashed box) found in the supermotifs (top) in other annotated SINE B1 (middle) and outside any annotated SINE B1 (bottom). The consensus sequence of the canonical SINE B1 is shown above the logos.

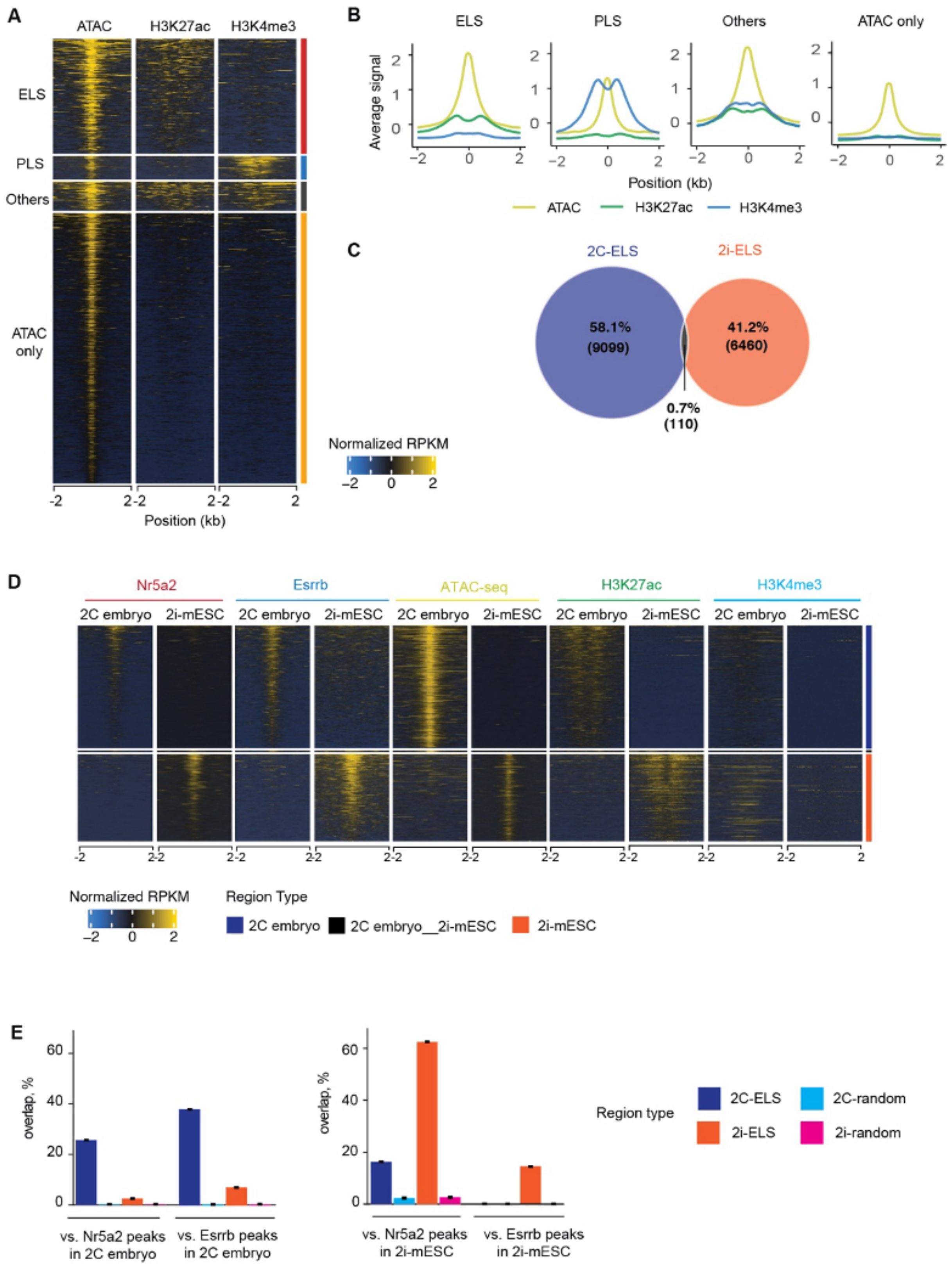


**Figure S4. Classification of cREs and cell-type specific ELS**
(**A and B**) Classification of cREs by epigenetic signatures in 2-cell embryos. Heatmaps (A) and line plots (B) are shown with ATAC-seq, H3K27ac and H3K4me3 enrichments (Z-score normalized). (**C**) Eular diagram showing overlap between 2C-ELS and 2i-ELS. (**D**) Heatmaps shows classified 2-cell embryo specific, 2i-mESC specific or common ELS with Nr5a2, Esrrb,

ATAC-seq, H3K27ac and H3K4me3 enrichments (*Z*-score normalized). (**E**) Bar charts showing the percentage of ELS from both cell types that are overlapped with Nr5a2 or Esrrb peaks in 2-cell embryos and 2i-mESCs.

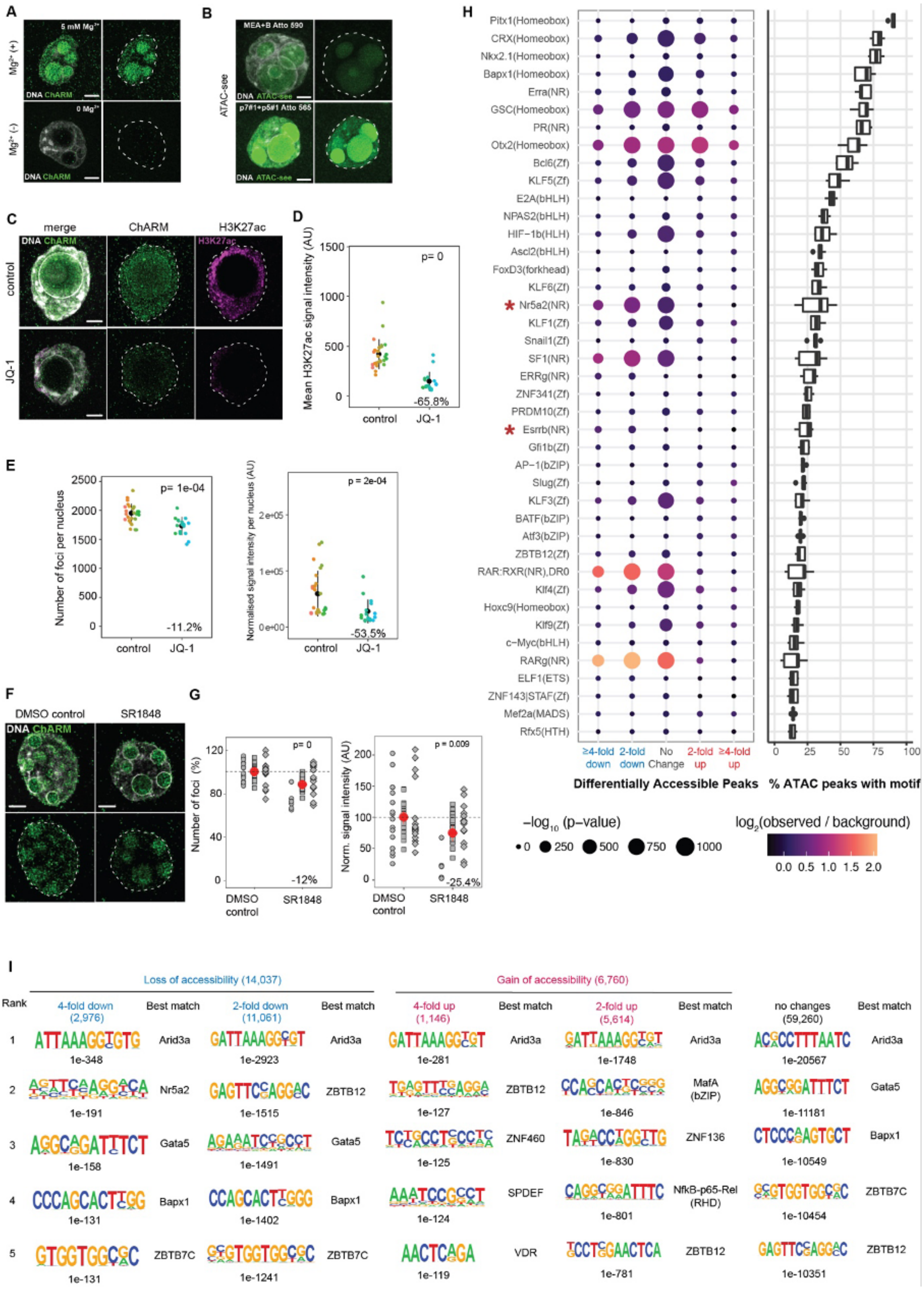

A
5 mM Mg2+
Mg2+ (+)
0 Mg2+
Mg2+ (-)
DNA ChARM
B
ATAC-see
MEA+B Atto 590
p7#1+p5#1 Atto 565
DNA ATAC-see
C
merge
ChARM
H3K27ac
control
JQ-1
D
Mean H3K27ac signal intensity (AU)
p= 0
-65.8%
control
JQ-1
E
Number of foci per nucleus
p= 1e-04
-11.2%
Normalised signal intensity per nucleus (AU)
p = 2e-04
-53.5%
control
JQ-1
F
DMSO control
SR1848
G
Number of foci (%)
p= 0
-12%
Norm. signal intensity (AU)
p = 0.009
-25.4%
DMSO control
SR1848
H
Pitx1(Homeobox)
CRX(Homeobox)
Nkx2.1(Homeobox)
Bapx1(Homeobox)
Erra(NR)
GSC(Homeobox)
PR(NR)
Otx2(Homeobox)
Bcl6(Zf)
KLF5(Zf)
E2A(bHLH)
NPAS2(bHLH)
HIF−1b(HLH)
Ascl2(bHLH)
FoxD3(forkhead)
KLF6(Zf)
Nr5a2(NR)
KLF1(Zf)
Snail1(Zf)
SF1(NR)
ERRg(NR)
ZNF341(Zf)
PRDM10(Zf)
Esrrb(NR)
Gfi1b(Zf)
AP−1(bZIP)
Slug(Zf)
KLF3(Zf)
BATF(bZIP)
Atf3(bZIP)
ZBTB12(Zf)
RAR:RXR(NR),DR0
Klf4(Zf)
Hoxc9(Homeobox)
Klf9(Zf)
c−Myc(bHLH)
RARg(NR)
ELF1(ETS)
ZNF143|STAF(Zf)
Mef2a(MADS)
Rfx5(HTH)
≥4-fold down
2-fold down
No Change
2-fold up
≥4-fold up
Differentially Accessible Peaks
% ATAC peaks with motif
−log10 (p−value)
log2(observed / background)
I
Loss of accessibility (14,037)
Gain of accessibility (6,760)
Rank
4-fold down (2,976)
Best match
2-fold down (11,061)
4-fold up (1,146)
2-fold up (5,614)
no changes (59,260)
1e-348 Arid3a
1e-2923 Arid3a
1e-281 Arid3a
1e-1748 Arid3a
1e-20567 Arid3a
1e-191 Nr5a2
1e-1515 ZBTB12
1e-127 ZBTB12
1e-846 MafA (bZIP)
1e-11181 Gata5
1e-158 Gata5
1e-1491 Gata5
1e-125 ZNF460
1e-830 ZNF136
1e-10549 Bapx1
1e-131 Bapx1
1e-1402 Bapx1
1e-124 SPDEF
1e-801 NfkB-p65-Rel (RHD)
1e-10454 ZBTB7C
1e-131 ZBTB7C
1e-1241 ZBTB7C
1e-119 VDR
1e-781 ZBTB12
1e-10351 ZBTB12

**Figure S5. Validation of ChARM and Omni ATAC-seq**
(**A**) Representative images showing $Mg^{2+}$-dependent tagmentation. Scale bars are 5 μm. (**B**) Representative images of ATAC-see with the original fluorescent DNA adapter (top) and fluorescent conjugates of the sci-ATAC-seq adapters used for ChARM (bottom). Scale bars are 5 μm. (**C**) Representative images showing ChARM in 2-cell embryos treated with BRD4 inhibitor JQ-1 (1.2 μM). Scale bars are 5 μm. (**D and E**) Scatterplot shows H3K27ac signal intensity (D) and the number of foci and normalized signal intensity of ChARM (E). The measurement per nucleus is normalised to the mean of the corresponding control. P-value of pairwise non-parametric Mann Whitney U test between pooled experimental and control distributions is shown. Sample sizes are: control: n = 26; JQ-1: n = 17 nuclei. Scale bars are 5 μm. (**F**) Representative images showing ChARM in 2-cell embryos treated with Nr5a2 antagonist SR1848 (10 μM). Scale bars are 5 μm. (**G**) Scatterplot shows the relative percentage of the number of ChARM foci per nucleus in E. Each measurement per nucleus from replicate experiments is normalized to the mean of the corresponding control and plotted in different shapes. The average values are shown as red dots. P-value of pairwise non-parametric Mann Whitney U test between pooled experimental and control distributions is shown. Sample sizes are: control: n = 17, 27, 18; SR1848: n = 6, 20, 14 nuclei. Scale bars are 5 mm. (**H**) Enrichment of known transcription factor motifs in differentially accessible ATAC peaks upon SR1848 treatment. The dot size corresponds to the -log10 P-value of the enrichment. Color scale indicates the log2 ratio of the proportion of peaks that contain motifs between the real sequence (observed) and shuffled sequence (background). Rows are ordered based on the proportion of peaks that contain the motif (boxplot). Asterisks indicate Nr5a2 and Esrrb. (**I**) *De novo* motif analysis showing top 5 motifs enriched in each different chromatin accessibility region. Enrichment p-value of sequences are noted.

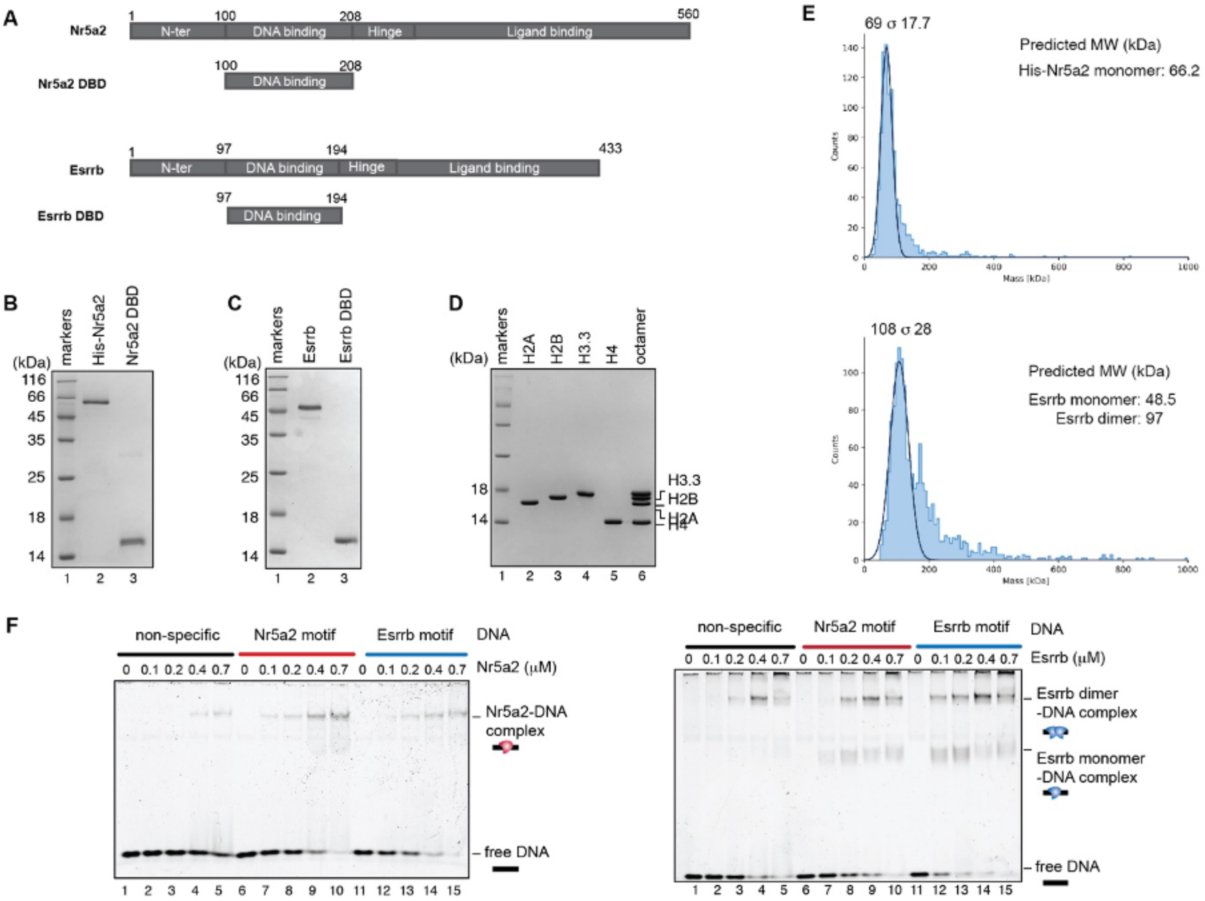


G

**Dissociation constants of recombinant Nr5a2 and Esrrb**

| Apparent $K_d$ (nM) | Nr5a2 | Nr5a2 DBD | Esrrb | Esrrb DBD |
|---|---|---|---|---|
| Non-specific | ND. * | – | ND.* | – |
| Nr5a2 motif | 5.62 ± 1* | 11.4 ± 4.87** | 625.53 ± 133* | – |
| Esrrb motif | 6.49 ± 0.35* | – | 8.56 ± 0.69* | 13.8 ± 3.4** |

*Apparent dissociation constants (KD) were derived from quantified FP measurements

**Apparent dissociation constants (KD) were derived from quantified EMSA bands

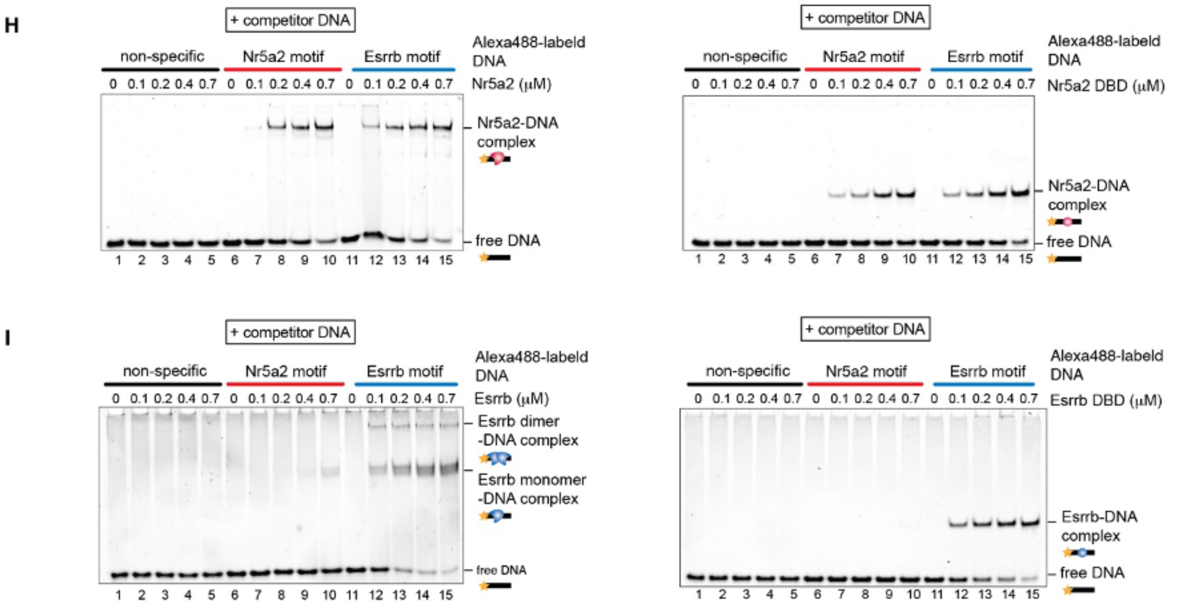

**Figure S6. Preparation of proteins and DNA-binding activity of Nr5a2 and Esrrb**
(**A**) *Mus musculus* Nr5a2 and Esrrb and their DNA-binding domains (DBDs) used in this study. (**B**) Purification of Nr5a2 full length and Nr5a2 DBD. Purified Nr5a2 (lane 2, 0.75 µg) and Nr5a2 DBD (lane 3, 0.75 µg) were analyzed by 18% SDS-PAGE. Lane 1 indicates molecular markers. (**C**) Purification of Esrrb full length and Esrrb DBD. Purified Esrrb (lane 2, 0.75 µg) and Esrrb DBD (lane 3, 0.75 µg) were analyzed by 18% SDS-PAGE. Lane 1 indicates molecular markers. (**D**) Purification of mouse histones and histone octamers. Lane 1 indicates molecular markers. Lanes 2-5 indicates purified H2A, H2B, H3.3 and H4, respectively. Lane 6 indicates the reconstituted histone octamers. The samples were analyzed by 18% SDS-PAGE. (**E**) Mass photometry profiles for Nr5a2 and Esrrb with determined average molecular mass (MS). The theoretical MS for Nr5a2 monomer including His-tag is 66.2 kDa. The theoretical MS for Esrrb monomer and dimer are 48.5 and 97 kDa, respectively. (**F**) DNA-binding specificity of Nr5a2 (left) and Esrrb (right) in the absence of competitor DNA. Three independent experiments were performed, and the reproducibility was confirmed. (**G**) Apparent dissociation constants ($K_d$) values measured by fluorescence polarization assay or EMSA. (**H and I**) EMSAs in the presence of competitor DNA with Nr5a2 (H) or Essrb (I). Lanes 1-5, lanes 6-10 and lanes 11-15 indicate results for the DNA-binding with 5'-Alexa488-labeled 25 bp dsDNA non-specific sequence, Nr5a2 motif or Esrrb motif, respectively. Two independent experiments were performed, and the reproducibility was confirmed.

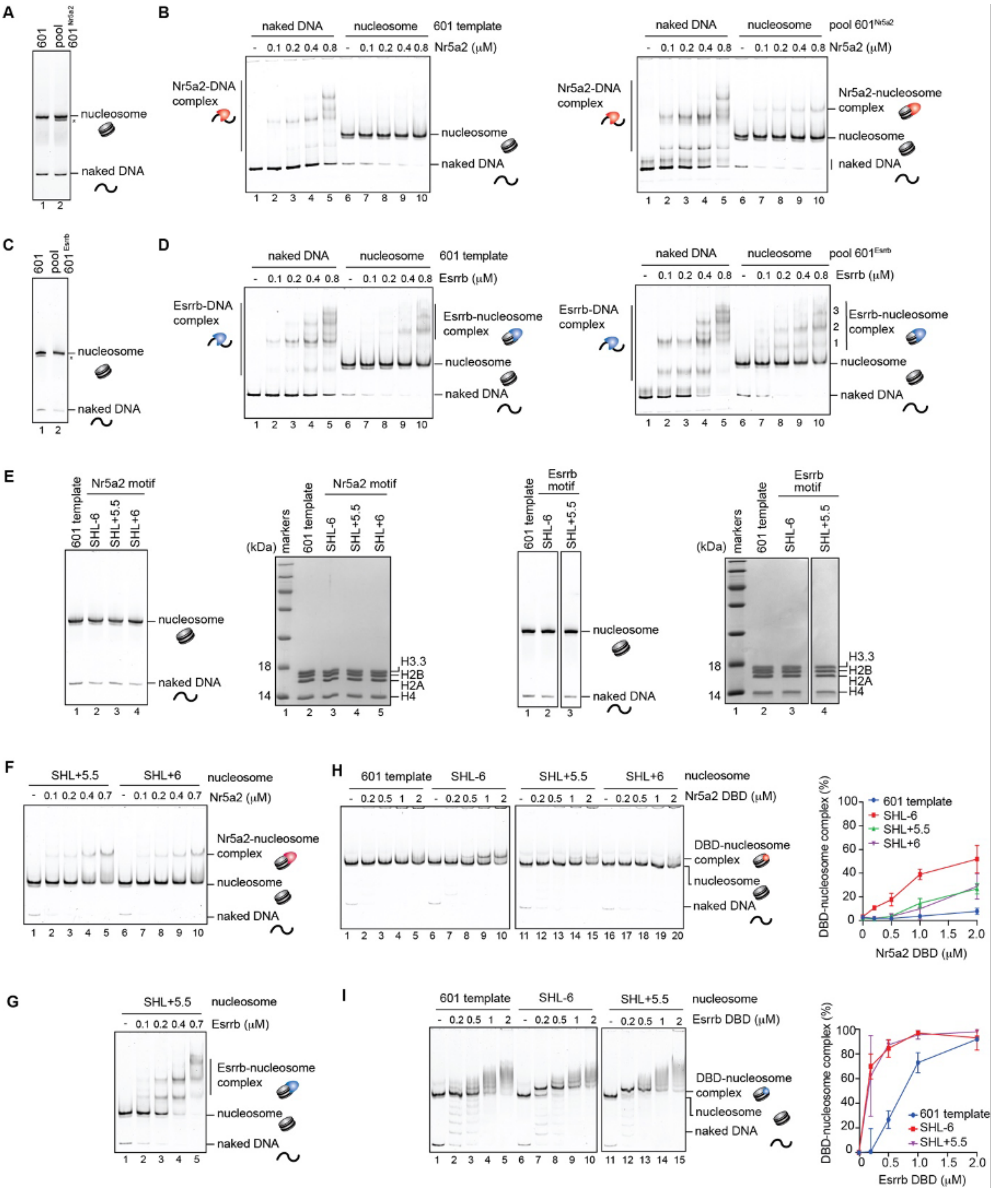


### Figure S7. Nucleosome binding activity of Nr5a2 and Esrrb

(**A**) Purified 601 template nucleosome (lane 1) and nucleosome libraries containing Nr5a2 motif (pool 601$^{Nr5a2}$, lane 2) were analyzed by native-PAGE with SYBR safe staining. Asterisk indicates hexasome which lacks a histone H2A-H2B dimer. (**B**) EMSA of 601 template or pool 601$^{Nr5a2}$ with Nr5a2. The naked DNA or nucleosome (100 nM) was mixed with Nr5a2 (0, 0.1, 0.2, 0.4 and 0.8 μM). lanes 1-5 and lanes 6-10 indicate results with naked DNA and nucleosome, respectively. One band corresponding to the Nr5a2-nucleosome complexes was cut and purified as TF-bound fraction. (**C**) Purified 601 template nucleosome (lane 1) and nucleosome libraries containing Esrrb motif (pool 601$^{Esrrb}$, lane 2) were analyzed by native-PAGE with SYBR safe staining. Asterisk indicates hexasome. (**D**) EMSA of 601 template or pool 601$_{Esrrb}$ with Esrrb. The naked DNA or nucleosome (100 nM) was mixed with Esrrb (0, 0.1, 0.2, 0.4 and 0.8 μM). lanes 1-5 and lanes 6-10 indicate results with naked DNA and nucleosome, respectively. Three bands corresponding to the Esrrb-nucleosome complexes were cut and purified as TF-bound fraction. (**E**) Purified nucleosomes containing Nr5a2 motif at SHL-6, SHL +5.5 and SHL+6 and Esrrb motif at SHL-6 and SHL+5.5 were analyzed by 6% native-PAGE with SYBR safe staining. The histone contents of the purified nucleosomes were analyzed by 18% SDS-PAGE. (**F**) Representative data of EMSA with SHL+5.5 and SHL+6 nucleosome. Quantification of the result are shown in Fig. 6E. (**G**) Representative data of EMSA with SHL+5.5 nucleosome. Quantification of the result are shown in Fig. 6F. (**H**) Nucleosome binding assay of Nr5a2 DBD. Nr5a2 DBD (0, 0.2, 0.5, 1 and 2 μM) was incubated with the nucleosome (50 nM), and the nucleosome and DBD-nucleosome complex were separated by 6% native-PAGE. Lanes 1-5, 6-10, 11-15 and 16-20 and indicate results for the nucleosome-binding experiments with the 601 template, SHL-6, SHL+5.5 and SHL+6, respectively. Quantification of the results is shown in right panel. The average values of three independent experiments are shown with the SD values. (**I**) Nucleosome binding assay of Esrrb DBD. Lanes 1-5, 6-10, 11-15 and 16-20 and indicate results for the nucleosome-binding experiments with the 601 template, SHL-6 and SHL+5.5, respectively. Quantification of the results is shown in right panel. The average values of three independent experiments are shown with the SD values.
